# Intrinsic checkpoint deficiency during cell cycle re-entry from quiescence

**DOI:** 10.1101/558783

**Authors:** Jacob Peter Matson, Amy M. House, Gavin D. Grant, Huaitong Wu, Joanna Perez, Jeanette Gowen Cook

## Abstract

The authors find that human cells re-entering the cell cycle from quiescence have both an impaired p53-dependent DNA replication origin licensing checkpoint and slow origin licensing. This combination makes every first S phase underlicensed and hypersensitive to replication stress.

**ABSTRACT:** To maintain tissue homeostasis, cells transition between cell cycle quiescence and proliferation. An essential G1 process is Minichromosome Maintenance complex (MCM) loading at DNA replication origins to prepare for S phase, known as origin licensing. A p53-dependent origin licensing checkpoint normally ensures sufficient MCM loading prior to S phase entry. We used quantitative flow cytometry and live cell imaging to compare MCM loading during the long first G1 upon cell cycle entry and the shorter G1 phases in the second and subsequent cycles. We discovered that despite the longer G1 phase, the first G1 after cell cycle re-entry is significantly underlicensed. As a result, the first S phase cells are hypersensitive to replication stress. This underlicensing is from a combination of slow MCM loading with a severely compromised origin licensing checkpoint. The hypersensitivity to replication stress increases over repeated rounds of quiescence. Thus, underlicensing after cell cycle re-entry from quiescence distinguishes a higher risk cell cycle that promotes genome instability.

## INTRODUCTION

Proliferating mammalian cells initiate DNA replication at thousands of DNA replication origins every cell cycle. Replication origins are chromosomal loci where DNA synthesis initiates in S phase. The Minichromosome Maintenance complex (MCM) is an essential component of the helicase that unwinds DNA to initiate replication. MCM loading dynamics are tightly regulated and coordinated with cell cycle progression to achieve precise and complete genome duplication (Bell and Labib, 2016). Cells prepare for DNA replication in S phase by loading MCMs at replication origins in the preceding G1 phase, a process called “origin licensing.” The amount of DNA-loaded MCM increases as cells progress through G1 until reaching a maximum per cell at the G1/S transition (Siddiqui et al., 2013; Remus and Diffley, 2009). Once cells enter S phase, multiple overlapping mechanisms block any *new* MCM loading, thus restricting origin licensing activity to G1 phase (Truong and Wu, 2011; Arias and Walter, 2007). Cells block MCM loading outside of G1 phase to prevent genotoxic re-replication, which causes aneuploidy, double strand breaks, gene amplification, and genome instability (Neelsen et al., 2013; Truong and Wu, 2011; Arias and Walter, 2007). MCMs unwind DNA in S phase, travel with replication forks, and MCMs are unloaded throughout S phase as replication forks terminate (Maric et al., 2014; Moreno et al., 2014).

Cells may encounter replication stress and replication fork stalling during S phase from a variety of endogenous and exogenous sources. A stalled replication fork can be rescued if MCM at a nearby licensed origin initiates a new fork to replicate the intervening DNA (Alver et al., 2014; Yekezare et al., 2013). Since MCM loading is tightly restricted to G1 phase, cells typically load excess MCM to license 5-10 fold more origins than they would require to complete S phase if there were no replication stress at all. These excess licensed origins function as “dormant origins”, and are activated as needed to respond to local replication stress and rescue stalled forks (Woodward et al., 2006; Ge et al., 2007; Ibarra et al., 2008). Cells with considerably less loaded MCM can still complete a normal S phase (Ge et al., 2007). Nonetheless, if cells enter S phase with less loaded MCM they are “underlicensed” with fewer dormant origins, and are hypersensitive to replication stress. In addition, animal models illustrate the long-term consequences of underlicensing. Mice heterozygous for MCM null alleles or homozygous for hypomorphic MCM alleles have less MCM loading, increased replication stress, and defects in highly proliferative tissues such as hematopoietic cells and intestinal crypts (Alvarez et al., 2015; Pruitt et al., 2007). In addition, these mice are prone to genomic instability, premature aging, and cancer (Kunnev et al., 2010; Shima et al., 2007; Pruitt et al., 2007).

Since dormant origins are critical to protect cells from replication stress and its consequent genome instability, cells have a control mechanism to ensure sufficient origin licensing. An origin licensing checkpoint in G1 phase of untransformed mammalian cells ensures abundant licensing in G1 phase before S phase entry (Shreeram et al., 2002; Nevis et al., 2009; Liu et al., 2009; Gillespie et al., 2007). Artificially reducing MCM loading in checkpoint-proficient cells delays the late G1 activation of Cyclin E/Cyclin Dependent Kinase 2 (CDK2) (Nevis et al., 2009; McIntosh and Blow, 2012). Delayed Cyclin E/CDK2 activation delays the phosphorylation of multiple substrates that drive S phase entry (Giacinti and Giordano, 2006). Delaying CDK2 activity lengthens G1 phase and ensures that cells do not enter S phase with low levels of origin licensing. Moreover, this checkpoint is p53-dependent (Nevis et al., 2009; Liu et al., 2009; Shreeram et al., 2002), thus a common genetic perturbation in transformed cancer cells likely compromises the normal coordination of origin licensing and S phase onset.

Given the importance of coordinating G1 length with the progress of origin licensing for robust S phase completion, we considered natural circumstances where G1 length changes. One example is the short G1 phase of pluripotent stem cells. We previously found that stem cells load MCM faster than differentiated cells with long G1 phases to achieve the same amount of loaded MCM at S phase entry (Matson et al., 2017). An alternative example is the long G1 after cell cycle re-entry from quiescence (Coller, 2007). Cell cycle quiescence, or “G0,” is a reversible cell cycle exit to a nondividing state. G0 is distinct from a G1 arrest; it is an active state requiring upregulation of anti-apoptotic, anti-senescent, and anti-differentiation genes as well as repression of cell cycle genes (Coller et al., 2006; Sang et al., 2008; Litovchick et al., 2007; Sagot and Laporte, 2019). The longer G1 phase during re-entry likely reflects the need to reactivate and express genes repressed in G0 and other fundamental differences in G1 regulation.

The unique features of G0 and the first G1 phase suggests that origin licensing may be distinctly regulated during cell cycle re-entry. We entertained the notion that the increased G1 length may allow more time for MCM loading, causing a meaningful increase in DNA-loaded MCM when entering S phase compared to subsequent proliferating cell cycles. To test this idea, we used single cell flow cytometry to measure the amount of loaded MCM at S phase entry and live cell imaging to measure cell cycle timing to determine if the amount of loaded MCM differs between cell cycle re-entry and active proliferation. Surprisingly, we discovered instead that cells re-entering the cell cycle from G0 are in fact, routinely and significantly underlicensed rendering them hypersensitive to replication stress compared to proliferating cells. This finding is consistent with a recent report that the first S phase experiences higher spontaneous replication stress, though the source of that endogenous stress was not identified (Daigh et al., 2018). We report here that MCM loading is slow in the first G1 phase and furthermore, that the first cell cycle has a severely compromised origin licensing checkpoint relative to the robust checkpoint in actively proliferating cells. This combination promotes premature S phase entry and underlicensing. To our knowledge, these results demonstrate the first naturally underlicensed cell cycle and suggest that repeated cell cycle re-entry from G0 is particularly hazardous for long-term genome stability.

## RESULTS

### G0 cells re-entering the cell cycle are underlicensed compared to actively proliferating cells

Cells re-entering the cell cycle from G0 have a long G1 before S phase entry relative to actively proliferating cells (Coller, 2007). We hypothesized that the difference in G1 length and unique aspects of the G0 to G1 transition could alter the timing or amount of loaded MCM at S phase entry in normal human cells. The amount of MCM loaded at the onset of S phase determines replication stress tolerance and thus overall replication success. For this reason we focused on quantifying how much MCM is loaded at the onset of S phase (G1/S transition). Standard synchronization and immunoblotting is an inadequate approach however because G0 cells re-enter G1 phase semi-synchronously, particularly untransformed cells. Even in the first cycle after re-entry, there is no time point near the G1/S transition that is not a mix of different phases (e.g. G1 and S phase cells), and the following second cell cycle is nearly asynchronous (Wang et al., 2017; Kwon et al., 2017). We therefore used a previously published and validated single cell flow cytometry assay to measure DNA-loaded MCM at the G1/S transition (Matson et al., 2017; Håland et al., 2015; Moreno et al., 2016; Carroll et al., 2018; Hiraga et al., 2017). We examined proliferating Retinal Pigmented Epithelial cells (RPE1) whose only genetic alteration is the introduction of telomerase (RPE1-hTert). We extracted soluble proteins with nonionic detergent so that DNA-bound proteins were retained, then fixed and stained with anti-Mcm2 antibody as a marker for the whole MCM2-7 complex, DAPI to measure DNA content, and for the nucleotide analog 5-Ethynyl-2’-deoxyuridine (EdU) to measure DNA synthesis (Fig. 1A). We defined MCM-positive cells based on an antibody negative control; grey cells in these plots are below this background for detecting DNA-loaded MCM (MCM^DNA^ neg.). Fig. S1A shows the flow cytometry gating scheme. We marked cells blue for 2C DNA content (G1) that are negative for DNA synthesis but positive for DNA-loaded MCM (G1-MCM^DNA^ pos.). MCM loading is unidirectional in G1 phase, increasing in G1 until the maximum amount of loaded MCM at the G1/S transition (Kuipers et al., 2011). Loaded MCMs are very stable and are only unloaded in S phase as replication terminates (Moreno et al., 2014; Maric et al., 2014). We marked cells orange for S phase based on EdU incorporation plus DNA-loaded MCM (S-MCM^DNA^ pos.). In actively proliferating populations with robust licensing prior to S phase entry, we consistently observe that the majority of cells reach a similar maximum loaded MCM per cell in G1 (blue) before becoming EdU positive (orange) in very early S phase. In this way, we accurately measured loaded MCM in single cells at very early S phase even in fully asynchronous populations. (Fig. 1A “G1/S” marked by the arrow).

**Figure 1.**
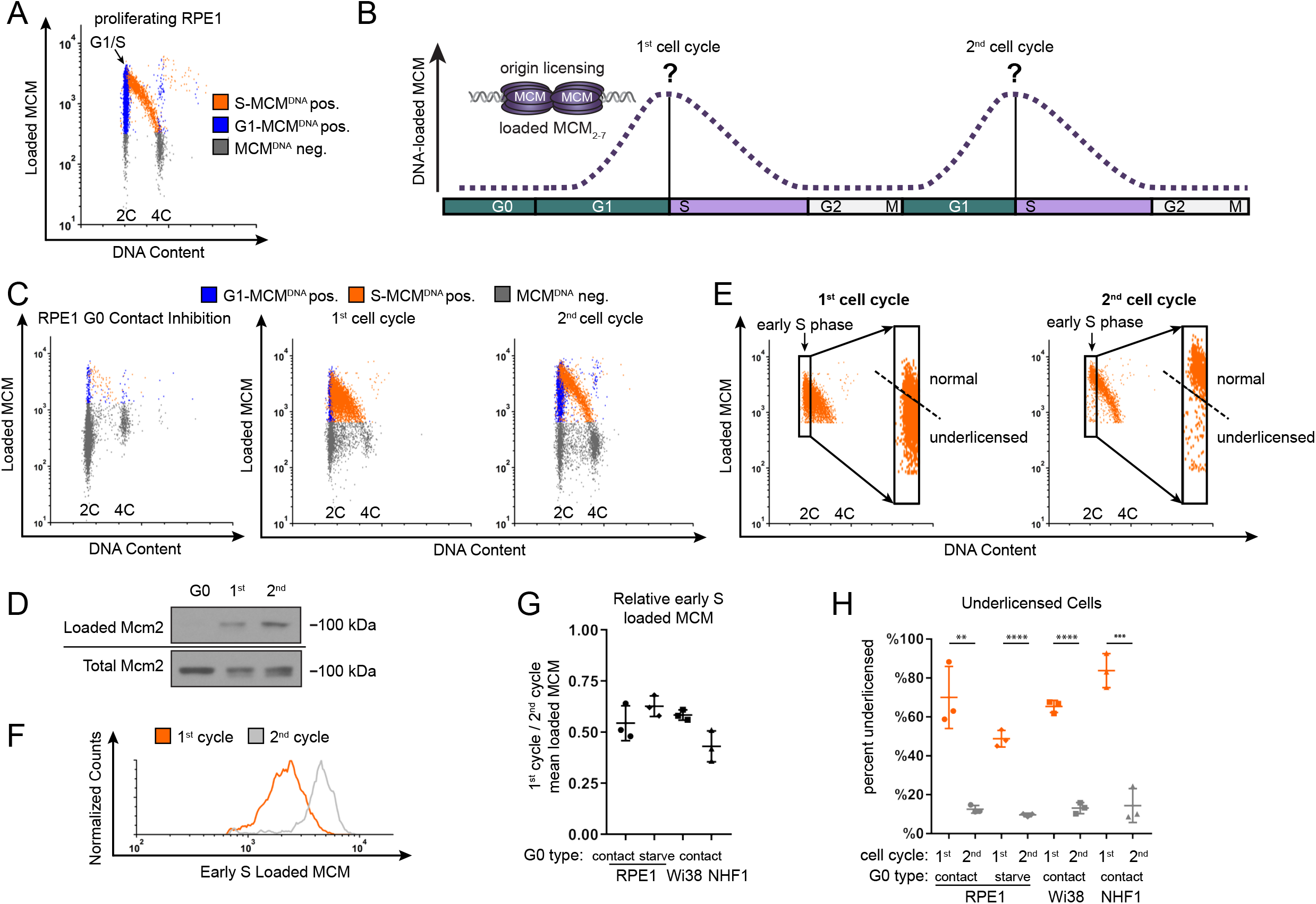
The first S phase after cell cycle re-entry from quiescence (G0) is underlicensed. **a**. Flow cytometry of proliferating RPE1-hTert cells extracted with nonionic detergent to measure DNA-bound protein (analytical flow cytometry for chromatin-bound proteins). Cells were labeled with 10 μM EdU for 30 minutes before harvesting then stained with anti-Mcm2 antibody to measure DNA loaded MCM, DAPI to measure DNA content, and EdU to measure DNA synthesis. Gates to define colors are provided in Fig. S1A: orange cells are S phase: EdU-positive + DNA-loaded MCM-positive, blue cells are G1 phase: EdU-negative + DNA-loaded MCM-positive, grey cells are all MCM-negative cells. Arrow indicates the G1/S transition. **b**. Cartoon of MCM loading in the first and second cell cycles. The amount of MCM loaded at the point of S phase entry determines the likelihood of replication success (indicated by question marks). At the start of this study, MCM loading before the first S phase after the long G1 relative to MCM loading in second and subsequent cell cycles was unknown. **c**. DNA-loaded MCM, DNA content (DAPI), and DNA synthesis (EdU) as in Fig. 1A. RPE1-hTert cells were synchronized in G0 by contact inhibition in the presence of growth serum and released from G0 into the cell cycle by replating. Cells were analyzed in G0, 24 hours after release (first cell cycle), and 48 hours after release (second cell cycle). **d**. Immunoblot of cells treated as in Fig. 1C, Samples were fractionated into DNA-loaded chromatin fractions and whole cell lysates then probed for MCM2. **e**. Gating of early S cells from Fig. 1C, showing only the S phase, MCM^DNA^-positive cells. Black rectangles define early S phase cells as MCM^DNA^-positive with 2C DNA content. Inset: Displays only early S phase cells. Cells above the dashed line were defined as normally licensed whereas cells below the line were underlicensed. **f**. Histogram measuring DNA-loaded MCM per cell in early S phase cells from Fig. 1E. The orange line is early S phase loaded MCM from first cell cycle, and the grey line is early S phase loaded MCM from second cell cycle. **g**. Comparison of early S phase DNA-loaded MCM presented as fold-change between the first and second cycles: RPE1-hTert synchronized by contact inhibition (e.g. Fig. 1E) by mitogen starvation (see also Fig. S1I, J), and Wi38, and NHF1-hTert G0 synchronized by contact inhibition (see also Fig. S1E-H). Comparisons are the mean intensity of DNA-loaded MCM in the first cell cycle divided by the corresponding mean in the second cell cycle. Horizontal bars indicate means, error bars mark standard deviation (SD), n=3 biological replicates. **h**. Percentage of underlicensed cells from first and second cell cycles in Fig. 1G. Horizontal bars indicate means, error bars mark standard deviation (SD), n=3 biological replicates. First and second cell cycles were compared by unpaired, two tailed t test. RPE1-hTert contact p=0.0035** RPE1 starve p<0.001****, Wi38 p<0.001****, NHF1 p=0.006***.

We then applied this method to measure how much MCM had been loaded by the time of the G1/S transition in the first cell cycle compared to the amount loaded at G1/S in the second cell cycle (Fig. 1B). We arrested RPE1-hTert cells in G0 by contact inhibition (in the presence of growth serum) for 48 hours. Contact-inhibited G0 cells showed a robust cell cycle exit to G0 with very little loaded MCM; 94% of cells were G0/G1 but only 1% were still in S phase (Fig. 1C and Fig. S1B). We then released cells to re-enter the cell cycle by plating at sub-confluent cell density. G0 cells also expressed the expected high levels of p27 and low Cyclin D1 compared to cells in the first cycle (Fig. S1C). We pulse-labeled cells with EdU and harvested some cells in G0 or at 24 hours when they were a mix of first G1 and ~50-70% first S phase cells. In a separate continuous EdU labeling experiment in which EdU was added at the time of release, we determined that nearly 100% of cells had entered S phase by 28 hours (Fig. S1D), and almost no cells remained in G0/G1. Thus, we concluded that additional cells we harvested 48 hours after release had completed their first S phase and were into their second cell cycle.

Strikingly, the amount of loaded MCM at the G1/S transition was markedly different between the first and second cell cycles after G0. Cells in the first cell cycle progressed into S phase (orange cells) with substantially less MCM loaded than cells in the second cell cycle. By the second cell cycle, G1 cells progressed into S phase as a tight group with relatively high amounts of loaded MCM (transition from blue to orange). In contrast, many cells in the first cell cycle entered S phase with low amounts of loaded MCM. This behavior creates the filled orange triangle on flow cytometry plots of the first S phase instead of the normally clear region under a high arc characteristic of the second S phase. We also observed decreased loaded MCM in the first cycle using biochemical chromatin fractionation and immunoblotting (Fig. 1D).

To quantify the amount of loaded MCM at the G1/S transition, we analyzed the S phase cells from Fig. 1C, and defined very early S phase cells as EdU-positive with ~2C DNA content (Fig. 1E black rectangles). Early S phase cells in the first cycle had a broad range of loaded MCM levels that included many cells with low MCM, whereas early S phase cells in the second cycle primarily had high loaded MCM levels. The second cell cycle was nearly indistinguishable from asynchronously proliferating cells that had not been arrested. We thus used the second cell cycle to define the normal licensing levels, and classified cells with less MCM loaded in early S as “underlicensed” (Fig. 1E dashed line). To visualize and directly compare MCM loading distribution in different populations, we generated histograms of loaded MCM levels in early S phase cells (Fig. 1F). On this plot the differences between the first and second S phases were also readily apparent (compare orange and grey lines). We quantified the differences as fold change between the mean MCM loaded in the first and second S phases. Cells in the first cycle after re-entry from G0 loaded only half as much MCM as cells in the second cell cycle (Fig. 1G). We then used this same comparison to test if this underlicensed first cell cycle is common among different untransformed cell lines or methods of quiescence induction. We observed similar underlicensing after quiescence induction in RPE1-hTert by serum starvation and re-stimulation and in two normal fibroblast cell lines arrested by contact inhibition and release (Fig. 1G and Fig. S1E-J). We observed that not only was the mean MCM loaded by the first S phase half that of subsequent S phases, but also that the majority of first S phase cells were underlicensed (Fig. 1H). Thus, re-entry from G0 is characterized by substantial underlicensing in the first cell cycle.

### Cells re-entering S phase from G0 are hypersensitive to replication stress

Cells typically load extra MCM in G1 to license dormant origins so they can tolerate replication stress in S phase (Woodward et al., 2006; Ge et al., 2007; Ibarra et al., 2008). Given our observations that the first S phase is underlicensed relative to the subsequent S phase, we hypothesized that cells re-entering the cell cycle from G0 would be hypersensitive to replication stress in the first S phase. To test that idea directly, we treated cells in the first or second cell cycle after G0 with low-dose gemcitabine, a drug that depletes nucleotides to cause replication stress (Fig. 2A). We used flow cytometry to measure the expression of γH2AX, a common replication stress marker (Ewald et al., 2007)(Fig. 2B, and Fig. S2A). We specifically analyzed mid-S phase cells to account for differences in cell cycle distribution, and we scored the number of γH2AX-positive S phase cells with expression equal to or greater than the top 5% of untreated cells (Fig. 2C, dashed line). By this measurement, cells in the first S phase after G0 are significantly more sensitive to replication stress than cells in the second S phase (Fig. 2D). Moreover, gemcitabine-treated first S phase cells expressed double the amount of γH2AX per cell than cells in the second S phase, suggesting that not only were more total cells exhibiting a replication stress response but also that there was more replication stress per cell (Fig. 2E, F). This hypersensitivity to replication stress in the underlicensed first cell cycle suggests that cell cycle re-entry is an inherently higher-risk cycle with respect to genome stability compared to subsequent cell cycles.

**Figure 2.**
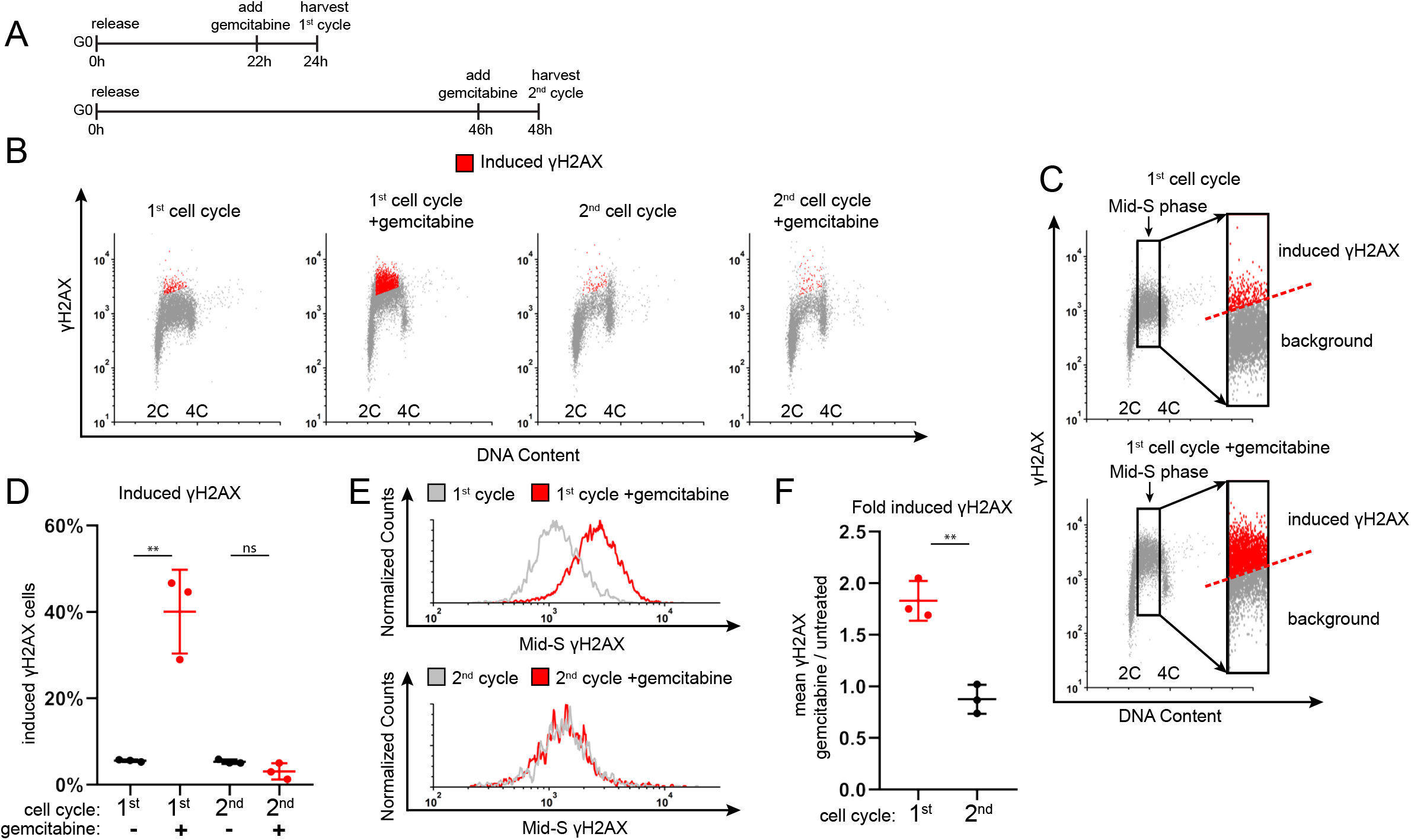
The first S phase after G0 is hypersensitive to replication stress. **a**. Diagram of experimental workflow. RPE1-hTert cells were synchronized in G0 by contact inhibition and released into the first cell cycle (24 hours) or second cell cycle (48 hours). Cells were treated with 50 nM gemcitabine or vehicle for the last 2 hours before harvesting for flow cytometry. **b**. Flow cytometry of chromatin-bound proteins in cells treated as in Fig. 2A and stained for DNA content (DAPI), and γH2AX (anti-H2AX phospho S139). Red cells are replication stress-induced γH2AX-positive (gemcitabine), as indicated in Fig. 2C. See also Fig. S2 for DNA-loaded MCM in these same cells. **c**. Gating of cells from Fig. 2B to define the threshold of γH2AX signal induced by gemcitabine. The rectangle isolates mid-S phase cells. Inset: Mid-S phase cells in which replication stress-induced γH2AX-positive cells (red) are defined as those equal to or above the top 5% of the corresponding untreated cells; these cells were also marked red in Fig. 2B. **d**. Percentage of replication stress-induced γH2AX in the first and second cell cycles from Fig. 2B. Horizontal bars indicate means, error bars mark standard deviation (SD), n=3 biological replicates. Untreated and gemcitabine-treated cells were compared by unpaired, two tailed t test. First cell cycle p=0.0035**; second cell cycle p=0.1174 (ns). **e**. Histograms of mid-S phase γH2AX intensity per cell from Fig. 2B; the upper panel is the first cell cycle and the lower panel is the second cell cycle. Grey lines are untreated cells, red lines are gemcitabine-treated cells. **f**. Comparison of γH2AX intensity per cell presented as fold-change between gemcitabine-treated and untreated cells from cells in Fig. 2E. Horizontal bars indicate means, error bars mark standard deviation (SD), n=3 biological replicates. First and second cell cycles were compared by unpaired, two tailed t test, p=0.0023**.

Many cells *in vivo* switch between periods of active proliferation and periods of G0, repeatedly re-entering the cell cycle to proceed through this presumably underlicensed first cell cycle. We considered the possibility that repeated cell cycle exit and re-entry would lead to additional replication stress sensitivity over time. To measure accumulation of replication stress during repeated cell cycle re-entry, we synchronized RPE1 cells in G0 by contact inhibition for 48 hours and released them into the cell cycle for about 48 hours (~2-3 cycles) and then they became contact inhibited again. We reiterated this procedure for a total of 3 rounds of G0 arrest and re-entry. In the third release, we treated with gemcitabine and measured the induction of γH2AX as before (Fig. S2B, C). Cells re-entering S phase from three repeated arrests in G0 (“3×G0”) were more likely to induce γH2AX cells than cells that been arrested only once (“1×G0”), and these cells were also more intensely γH2AX positive per cell (Fig. S2D, E). The increased replication stress sensitivity after three rounds of G0 varied somewhat due to inherent variations in the G0 synchronization over multiple rounds (Kwon et al., 2017; Wang et al., 2017). This variability was reflected by unpaired t test p values of ~p=0.15 for both the number of γH2AX positive cells and the γH2AX intensity Fig. S2D, E. Nevertheless, we observed a clear trend that repeated cell cycle re-entry from G0 increased sensitivity to replication stress.

### Proliferating epithelial cells have a robust p53-dependent origin licensing checkpoint

Our findings raised a larger question about the relationship between origin licensing and S phase entry. The consistently high amount of loaded MCM we observed prior to G1/S in the second and subsequent proliferating cell cycles is consistent with an active origin licensing checkpoint (e.g. Fig. 1A arrow). The origin licensing checkpoint is thought to delay S phase entry when the amount of loaded MCM is still low in untransformed cells by delaying the activation of G1 Cyclin Dependent Kinases (CDK) (McIntosh and Blow, 2012). Evidence for this checkpoint is low CDK activity and lengthening of G1 in proliferating cells treated to reduce MCM loading by RNAi-mediated depletion of MCM loading factors or by overproduction of the MCM loading inhibitor, Geminin (Shreeram et al., 2002; Liu et al., 2009; Nevis et al., 2009). The origin licensing checkpoint ensures that S phase begins with abundant licensed dormant origins to tolerate replication stress. Because we observed routinely underlicensed cells in the first S phase after G0 and consistently higher licensing in the second cell cycle, we hypothesized that the origin licensing checkpoint is primarily active in only the second and subsequent cell cycles.

Origin licensing checkpoint activity is cell type-dependent however (Shreeram et al., 2002). We therefore sought first to assess checkpoint status in proliferating untransformed epithelial cells (RPE1-hTert). We decreased origin licensing with siRNA targeting Cdc10 dependent transcript 1 (Cdt1), an essential MCM loading protein (Pozo and Cook, 2016). We used either a pool of four siCdt1 sequences (siCdt1 A) or a single independent siCdt1 (siCdt1 B) and analyzed Cdt1 protein by immunoblotting (Fig. 3B). We then tested both MCM loading and the length of G1 phase. Cdt1 depletion induced both a reduction in the rate of G1 phase MCM loading (Fig. S3A, B) and also a striking G1 lengthening. G1 length dramatically increased by several-fold (12 to 42 hours for siCdt1B), and the more profound Cdt1 depletion by siCdt1 B caused a greater G1 delay (Fig. 3F, green bars). Importantly, Cdt1 depletion alone changed neither the amount of early S loaded MCM (compare black and grey lines in Fig. 3C, and fold change in Fig. 3D) nor the percentage of underlicensed early S phase cells (Fig. 3E). By the time cells finally entered S phase, they had achieved normal amounts of loaded MCM. These observations indicate that actively proliferating RPE1 cells wait for the normal amount of loaded MCM in G1 phase before entering S phase.

**Figure 3.**
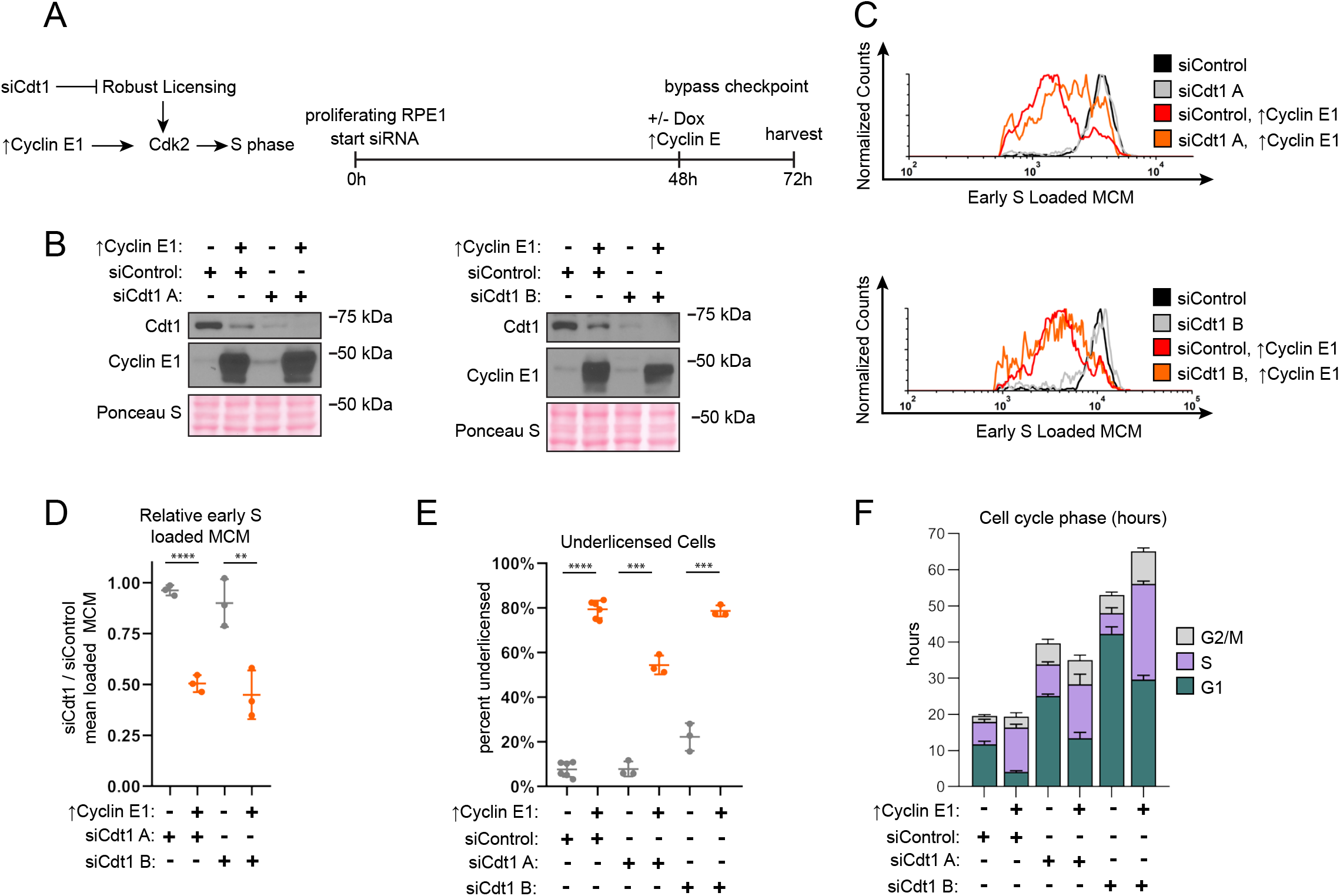
Cyclin E1 overproduction bypasses the licensing checkpoint in proliferating cells. **a**. Left: Model. Robust licensing promotes CDK2 activation and S phase entry. Cdt1 depletion by RNAi prevents robust licensing, CDK2 activation, and S phase entry. Cyclin E1 overproduction in Cdt1-depleted cells directly activates CDK2 independently of licensing and induces S phase entry. Right: Diagram of the experimental workflow. Proliferating RPE1-hTert cells containing integrated doxycycline inducible exogenous Cyclin E1 were treated with siControl, or independent siRNAs targeting Cdt1 (siCdt1 A, or siCdt1 B) for 72 hours. 100 ng/mL doxycycline to overproduce Cyclin E1 were added at 48 hours. **b**. Immunoblots of the indicated proteins in total protein lysates of cells treated as in Fig. 3A. ↑Cyclin E1 indicates addition of 100 ng/mL doxycycline at 48 hours to overproduce Cyclin E1. **c**. Loaded MCM in early S phase determined by flow cytometric analysis of cells treated as in Fig. 3A, measuring DNA content (DAPI), DNA-Loaded MCM (anti-Mcm2) and DNA synthesis (EdU). Black lines are siControl treated cells, grey lines are siCdt1 A (upper) or siCdt1 B (lower), red lines are siControl plus overproduced Cyclin E1. Orange lines are siCdt1 A (upper) or siCdt1 B (lower) plus overproduced Cyclin E1. See Fig. S3A for complete flow cytometry plots. **d**. Comparison of early S phase DNA-loaded MCM per cell from Fig. 3C. Values plotted are the ratio of mean loaded MCM in cells treated with siCdt1 and cells treated with siControl. Horizontal bars indicate means, error bars mark standard deviation (SD), n=3 biological replicates. Control and Cyclin E-overproducing cells were compared by unpaired, two tailed t test, siCdt1 A p<0.001****. siCdt1 B p=0.0096**. **e**. Percentage of underlicensed cells from cells treated as Fig. 3A. Horizontal bars indicate means, error bars mark standard deviation (SD), n=3 biological replicates. Control and Cyclin E-overproducing cells were compared by unpaired, two tailed t test. SiControl p<0.001****, siCdt1 A p=0.001***, siCdt1 B p=0.001***. **f**. Mean lengths of G1 (green), S (purple), and G2/M (grey) phases in hours for cells treated as Fig. 3A. G1 lengths: siControl = 11.7 hrs, siControl + ↑Cyclin E1 = 4.1 hours, siCdt1 A = 25.1 hrs, siCdt1 A + ↑Cyclin E1= 13.4 hrs, siCdt1 B = 42.3 hrs, siCdt1 B + ↑Cyclin E1 = 29.6 hours. Error bars are SD, n=minimum 3 biological replicates. Cell cycle phase percentages from DNA Content vs DNA synthesis were multiplied by the doubling time in hours to calculate hours for each phase. Doubling time is the mean of 3 biological replicates.

A cell cycle checkpoint that delays progression to the next phase is distinct from a simple incapacity to proceed to the next phase because it is possible for genetic alterations to bypass a checkpoint and induce premature cell cycle progression (Hartwell and Weinert, 1989). The origin licensing checkpoint delays the activation of Cyclin E/CDK2 (Fig. 3A) (Nevis et al., 2009). To test if the G1 phase delay can be bypassed in Cdt1-depleted RPE1-hTert cells, we overproduced Cyclin E1 to prematurely activate CDK2. We used flow cytometry to measure DNA-loaded MCM, plotting a histogram of early S phase loaded MCM as before (Fig. 3C and Fig. S3A). As expected from prior reports, Cyclin E1 overproduction in control cells shortened G1 phase nearly three-fold (siControl vs ↑Cyclin E1, Fig. 3F, green bars) (Matson et al., 2017; Resnitzky et al., 1994). In addition Cyclin E1 overproduction alone induced underlicensing as measured by the amount of early S loaded MCM (compare black and red lines, Fig. 3C, and the percentage of underlicensed cells, Fig. 3E) (Ekholm-Reed et al., 2004; Matson et al., 2017). Strikingly, when we overproduced Cyclin E1 after Cdt1 depletion (Fig. 3A), early S phase cells became severely underlicensed, quantified by both MCM loaded per cell (compare grey and orange lines, Fig. 3C and fold change in Fig. 3D), and the percentage of underlicensed cells (Fig. 3E). Moreover, G1 length decreased even in the Cdt1-depleted cells (Fig. 3F). We previously determined that the apparent decease in Cdt1 protein upon Cyclin E1 overproduction is an indirect effect of cell cycle phase distribution because Cdt1 is stable in G1 and unstable in S phase (Matson et al., 2017). Since Cyclin E1 overproduction bypassed the strong G1 delay in control Cdt1-depleted cells and induced S phase entry with low amounts of loaded MCM, the G1 delay is caused by a *bona fide* origin licensing checkpoint.

Full origin licensing checkpoint activity in normal human fibroblasts requires the p53 tumor suppressor (Fig. 4A)(Nevis et al., 2009). To test if p53-deficient epithelial cells are more likely to enter S phase underlicensed, we compared isogenic WT and p53 homozygous null RPE1-hTert cells (gift of P. Jallepalli, Rodriguez-Rodriguez et al., 2018). We analyzed the effects of reduced origin licensing from Cdt1 depletion on both G1 length and early S phase licensing status. We first noted by immunoblotting that Cdt1 depletion induced accumulation of both p53 and the CDK2 inhibitor p21, the product of a p53-inducible gene, whereas p53 null cells lacked both p53 and detectable levels of p21 (Fig. 4B). The absence of p53 had little effect on either G1 phase MCM loading (Fig. S3C) or the amount of MCM loaded by early S phase in otherwise unperturbed cells (compare black and grey lines in Fig 4C, fold change in Fig. 4D, and percentage of underlicensed cells in Fig 4E). As before, Cdt1 depletion slowed MCM loading in WT cells and increased G1 length by several-fold (Fig. 4F, Fig. S3D). In contrast, Cdt1-depleted cells lacking p53 entered S phase with significantly less MCM loaded (compare red and orange lines in Fig. 4C, fold change in Fig. 4D, and percentage of underlicensed cells in Fig. 4E). Moreover, unlike p53 WT cells, Cdt1 depletion in p53 null cells did not cause G1 lengthening (Fig. 4F). Thus, loss of p53 cripples the origin licensing checkpoint in proliferating cells, allowing inappropriate premature S phase entry of underlicensed cells. We note that even in control cells that had not been depleted of Cdt1, the p53 null cells entered S phase at a slightly lower amount of loaded MCM on average compared to p53 WT cells (compare black and red lines in Fig. 4C). We also detected a modest increase in the percentage of underlicensed p53 null siControl cells (Fig. 4E). Overall we conclude that these untransformed epithelial cells utilize a p53-dependent checkpoint to couple the timing of S phase entry to the status of origin licensing.

**Figure 4.**
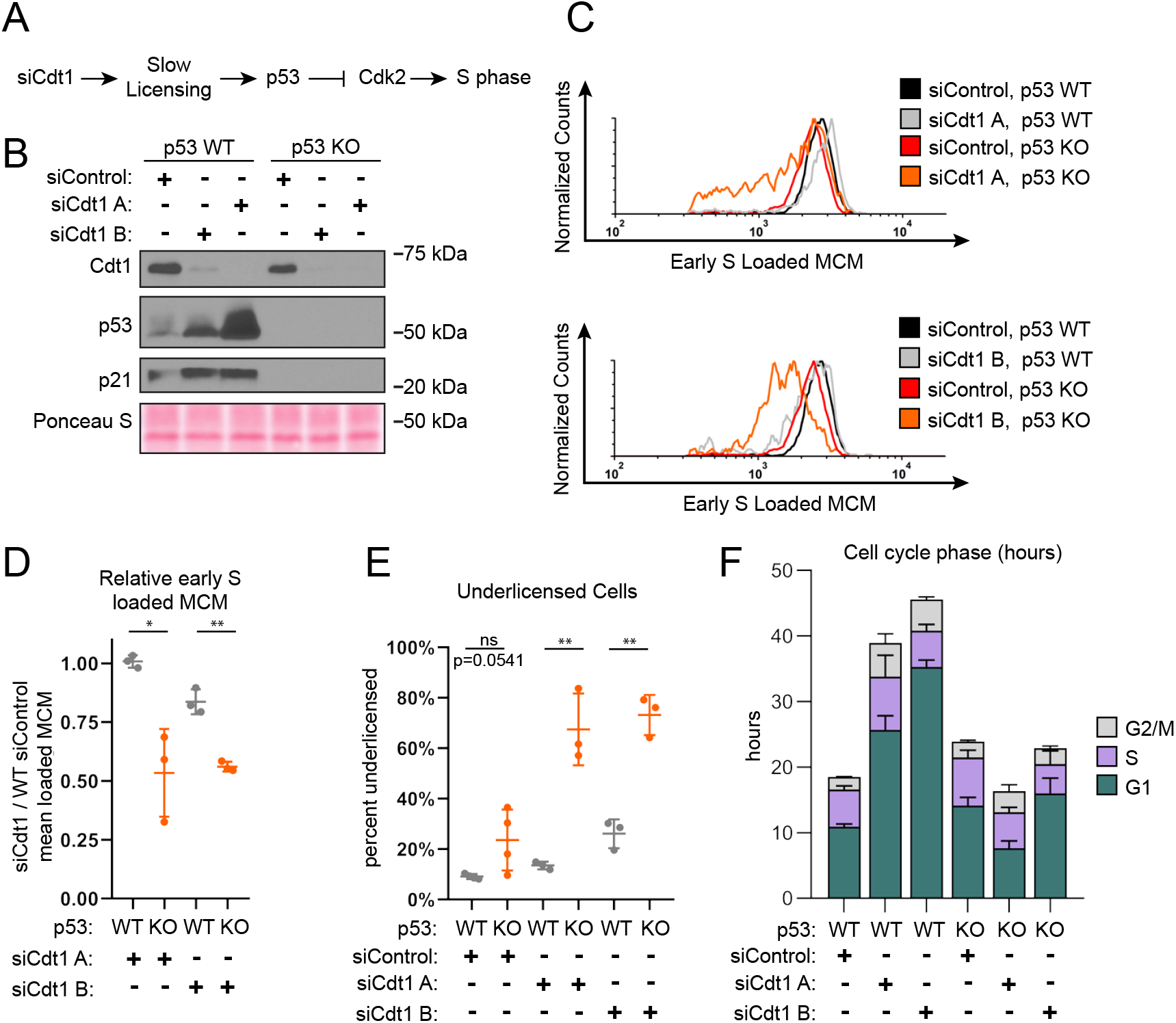
p53 loss cripples the licensing checkpoint and promotes underlicensing. **a**. Model. Cdt1 depletion causes slow origin licensing leading to p53-dependent inhibition of CDK2 which delays S phase entry. **b**. Immunoblot of the indicated endogenous proteins in total protein lysates of RPE1-hTert p53 WT or p53 null, (knockout “KO”) cells treated with siControl, siCdt1 A, or siCdt1 B for 72 hours. **c**. Loaded MCM in early S phase determined by flow cytometric analysis of cells treated as in Fig. 4B, measuring DNA Content (DAPI), loaded MCM (anti-Mcm2), and DNA synthesis (EdU). Histograms plot early S phase loaded MCM. Black lines are siControl + p53 WT, grey lines are siCdt1A + p53 WT (top) or siCdt1 B + p53 WT (bottom), red lines are siControl + p53 KO, orange lines are siCdt1 A + p53 KO (top) or siCdt1 B + p53 KO (bottom). The black siControl + p53 WT and red siControl + p53 KO are the same data on both histograms. Counts for all lines were normalized to siControl + p53 WT. **d**. Comparison of early S phase DNA-loaded MCM per cell from Fig. 4C. Values plotted are the ratio of mean loaded MCM in cells treated with siCdt1 and cells treated with siControl as indicated. Horizontal bars indicate means, error bars mark standard deviation (SD), n=3 biological replicates. WT and p53 KO cells were compared by unpaired, two tailed t test. SiCdt1 A p=0.0122*, siCdt1 B p=0.0011**. **e**. Percentage of underlicensed cells from Fig. 4C. Horizontal bars indicate means, error bars mark standard deviation (SD), n=3 biological replicates. WT and p53 KO cells were compared by unpaired, two tailed t test. SiControl p=0.0541 (ns), siCdt1 A p=0.0028**, siCdt1 B p=0.0011** **f**. Mean length of G1 (green), S (purple), and G2/M (grey) phases in hours for cells treated as Fig. 4C. G1 length: WT + siControl=10.9 hrs, WT + siCdt1 A = 25.7 hrs, WT + siCdt1 B= 35.5 hrs, p53 KO + siControl=14.1 hrs, p53 KO + siCdt1 A=7.6 hrs, p53 KO + siCdt1 B=16 hrs. Error bars are SD, n= minimum 3 biological replicates. Cell cycle phase percentages from DNA Content vs DNA synthesis were multiplied by the doubling time in hours to calculate hours for each phase. Doubling time is the mean of 3 biological replicates.

### The first G1 phase after G0 has an impaired origin licensing checkpoint

We had established in Figures 1 and 2 that cells re-entering the first cell cycle after G0 are routinely underlicensed relative to subsequent cycles. We hypothesized that the first cell cycle has an impaired origin licensing checkpoint that poorly couples the length of G1 phase to the status of MCM loading. To test that hypothesis, we compared actively proliferating cells treated with siCdt1 to G0 cells re-entering the first cycle also treated with siCdt1 (Fig. 5A). We measured cell cycle phase distribution by DNA Content (DAPI) and DNA synthesis (EdU) (Fig. 5B). The actively proliferating cells treated with siCdt1 increased the percentage of G1 cells as before, but cells re-entering the first cell cycle from G0 did not similarly respond to Cdt1 depletion (Fig. 5B, green bars). This difference is consistent with a weak origin licensing checkpoint in the first cell cycle. We also tested if the loss of p53 in the first cell cycle enhanced this apparent checkpoint deficiency (Fig. 5C). Unlike the proliferating cells in Fig. 4F, first cell cycle p53 null cells were no worse than first cell cycle WT cells (Fig. 5D). We note that siControl p53 null cells re-entering the first cycle also started S phase sooner than their corresponding p53 WT cells based on cell cycle distributions at the same time after G0 release (Fig. 5B 33% G1 vs, 16.6% G1 in Fig. 5D). The faster S phase entry by the p53 null cells could be due to both the impaired licensing checkpoint and the general loss of basal p21 protein (Fig. 5C) (Overton et al., 2014), among other possible p53-dependent effects on G1/S progression. The ability of actively proliferating cells to activate the licensing checkpoint to extend G1 combined with the observation that G0 cells do not extend G1 but instead enter S phase underlicensed strongly suggests that cells in the first cell cycle after G0 have an impaired origin licensing checkpoint.

**Figure 5.**
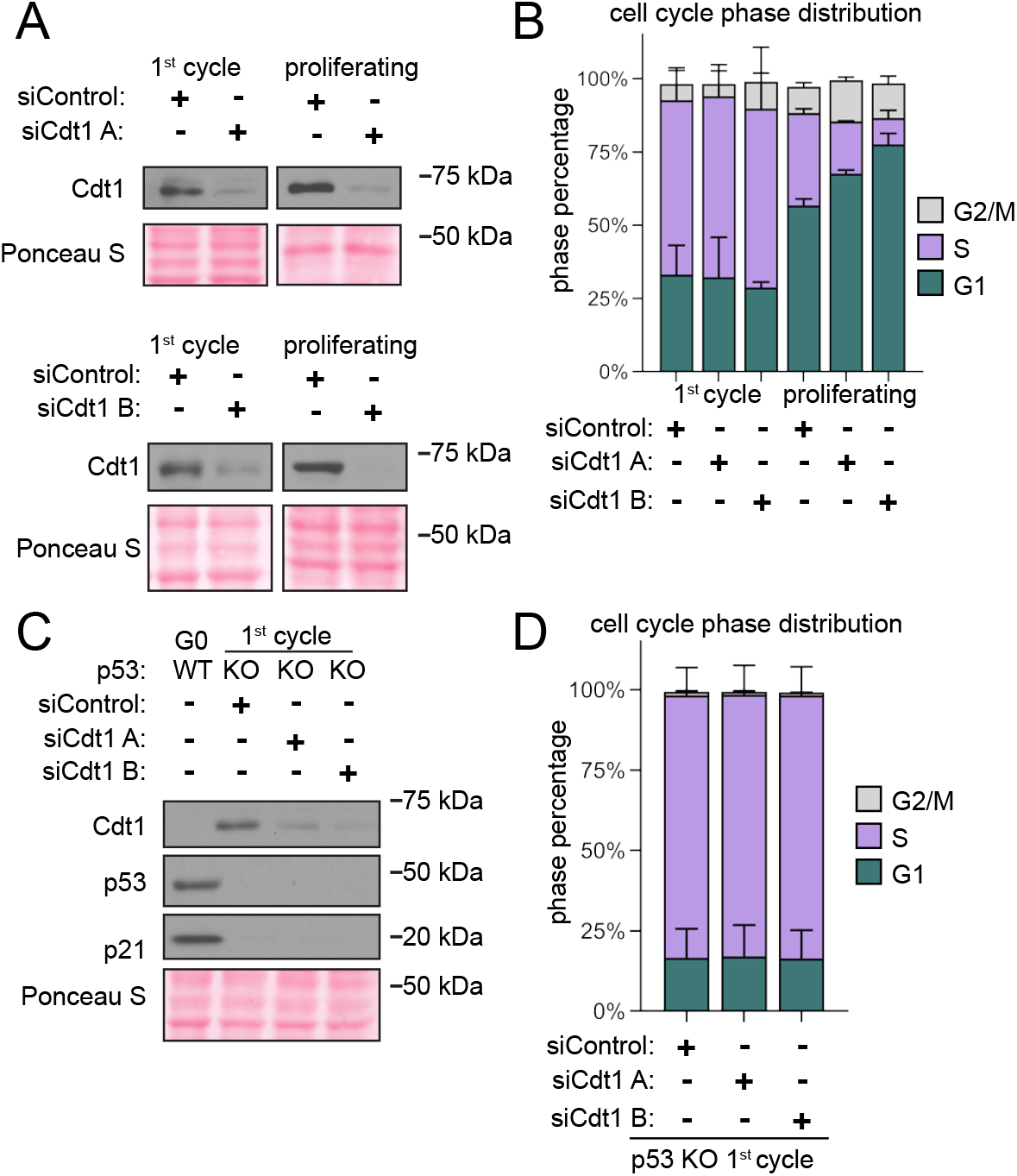
Cells re-entering the first G1 after G0 lack a licensing checkpoint-induced G1 delay. **a**. Immunoblot of total protein lysates of cells released from G0 into the first cell cycle with siControl or siCdt1 (24 hours after release) and of proliferating cells treated with siControl or siCdt1 for 72 hours. **b**. Cell cycle phase distribution of cells treated as Fig. 5A defined by DNA synthesis (EdU) and DNA Content (DAPI) using flow cytometry. Data plotted are mean percentage of G1 (green), S (purple), and G2/M (grey) phases. G1 percentage: 1^st^ cycle siControl = 33%, 1^st^ cycle siCdt1 A = 32.2%, 1^st^ cycle siCdt1 B = 28.7%, proliferating siControl = 56.7%, proliferating siCdt1 A = 67.5%, proliferating siCdt1 B = 77.5%. Error bars are SD, n= minimum 3 biological replicates. Cell cycle phase percentages from DNA Content vs DNA Synthesis. **c**. Immunoblot of total protein lysate of RPE1-hTert WT and p53 KO cells synchronized in G0 or released into the first cell cycle with siControl or siCdt1 (24 hours after release) as indicated. **d**. Cell cycle phase distribution of cells treated as Fig. 5C defined by DNA synthesis (EdU) and DNA Content (DAPI) using flow cytometry. Data plotted are mean percentage of G1 (green), S (purple), and G2/M (grey) phases. Error bars are SD, n= minimum 3 biological replicates. G1 percentage 16.6% (siControl) 17.0% (siCdt1 A) 16.4% (siCdt1B). Cell cycle phase percentages from DNA Content vs DNA Synthesis.

### G0 cells re-entering the first cell cycle load MCM to license origins slowly

We considered two explanations for underlicensing in the first S phase after G0: Cells may start loading MCM much later in the first G1 and, despite their longer G1, actually have less time to load than cells in the second cycle. Alternatively, cells in the first cycle may begin loading at the same time as in other cycles but they load MCM more slowly than cells in the second cycle. In both cases, the compromised origin licensing checkpoint is unable to extend G1 in response to the low MCM loading status resulting in an underlicensed first S phase. To distinguish between those explanations, we measured the nuclear accumulation of the Cell Division Cycle 6 (Cdc6) protein, an essential MCM loading protein. Cdc6 is degraded by the Anaphase promoting complex-Cdh1 (APC^Cdh1^) in G1 phase, both in cells re-entering the cell cycle and in proliferating cells (Petersen et al., 2000; Mailand and Diffley, 2005). CDK2/Cyclin E1 phosphorylates Cdc6 in late G1 to protect it from APC^Cdh1^, allowing Cdc6 protein to accumulate (Mailand and Diffley, 2005). Because Cdc6 is essential for MCM loading, Cdc6 accumulation is one of the limiting steps for MCM loading in G1. The time for MCM loading ends when S phase starts and Cdc6 is exported from the nucleus to the cytoplasm (Petersen et al., 1999) in addition to many other mechanisms that inhibit rereplication (Truong and Wu, 2011). Therefore, the time Cdc6 is detectable in nuclei is one proxy for the length of maximum available MCM loading time in G1 phase. Nuclear Cdc6 is also a marker for the functional border between G0/early G1 when MCM is not loaded (or loaded very slowly) and late G1 when MCM can be loaded.

We used live cell imaging of fluorescently tagged protein biosensors to compare the length of available MCM loading time between the first and second cell cycles after G0. We imaged an RPE1 cell line stably expressing three fluorescent fusion proteins: 1) full length Cdc6 fused to mVenus (Segev et al., 2016), 2) Proliferating Cell Nuclear Antigen (PCNA) fused to mTurq2 to track cell nuclei and the borders of S phase (Burgess et al., 2012; Grant, Kedziora et al., 2018), and 3) a CDK kinase activity sensor fused to mCherry (Hahn et al., 2009). This kinase sensor was previously established to report CDK2 activity in G1 and early S phase and both CDK2 and CDK1 activity in S and G2 phase; it is not a direct reporter of CDK4/6 activity (Spencer et al., 2013; Schwarz et al., 2018). CDK2 phosphorylates the reporter beginning in late G1 phase to induce export from the nucleus to the cytoplasm. The higher the ratio of cytoplasmic/nuclear signal, the higher the kinase activity. We imaged RPE1 cells expressing all three biosensors synchronized in G0 and released into the cell cycle for 72 hours, capturing an image every 10 minutes, starting 6.5 hours after release. PCNA-mTurq2 was present throughout the cell cycle and become punctate during S phase (Fig. 6A). Cdc6-mVenus was not present in G0 due to APC^Cdh1^-mediated degradation. Nuclear Cdc6 first appeared in G1 and increased until S phase at which point it was lost from the nuclei and accumulated in the cytoplasm instead. After mitosis, Cdc6 was degraded in early G1 phase, then nuclear Cdc6 increased again later in G1 (Fig. 6A). The CDK reporter (DHB-mCherry) was nuclear in G0 (low CDK activity) and gradually became cytoplasmic beginning in late G1 (high CDK activity)(Fig. 6A). We tracked 50 cells through the first and second cell cycles after G0 release. Fig. 6B shows an individual cell trace for nuclear Cdc6 in which the first full cell cycle took nearly 35 hours whereas the second cycle took only 20 hours mostly from the difference in G1 length. The time Cdc6 first appeared in G1 is marked “rise,” and the time of maximum nuclear Cdc6 in G1 is marked “peak” (Fig. 6B). Cells began S phase as measured by the first appearance of PCNA foci about 20-30 minutes after this peak (data not shown). The length of time Cdc6 was present in G1 is the time between nuclear Cdc6 rise and peak, and represents the “licensing window” for MCM loading in G1 (Fig. 6B). Cells reentering the first cell cycle after G0 had about double the licensing window of cells in the second cell cycle, and about twice as much time in G1 overall (Cdc6 peak time) as cells in the second cell cycle (Fig. 6C, D).

**Figure 6.**
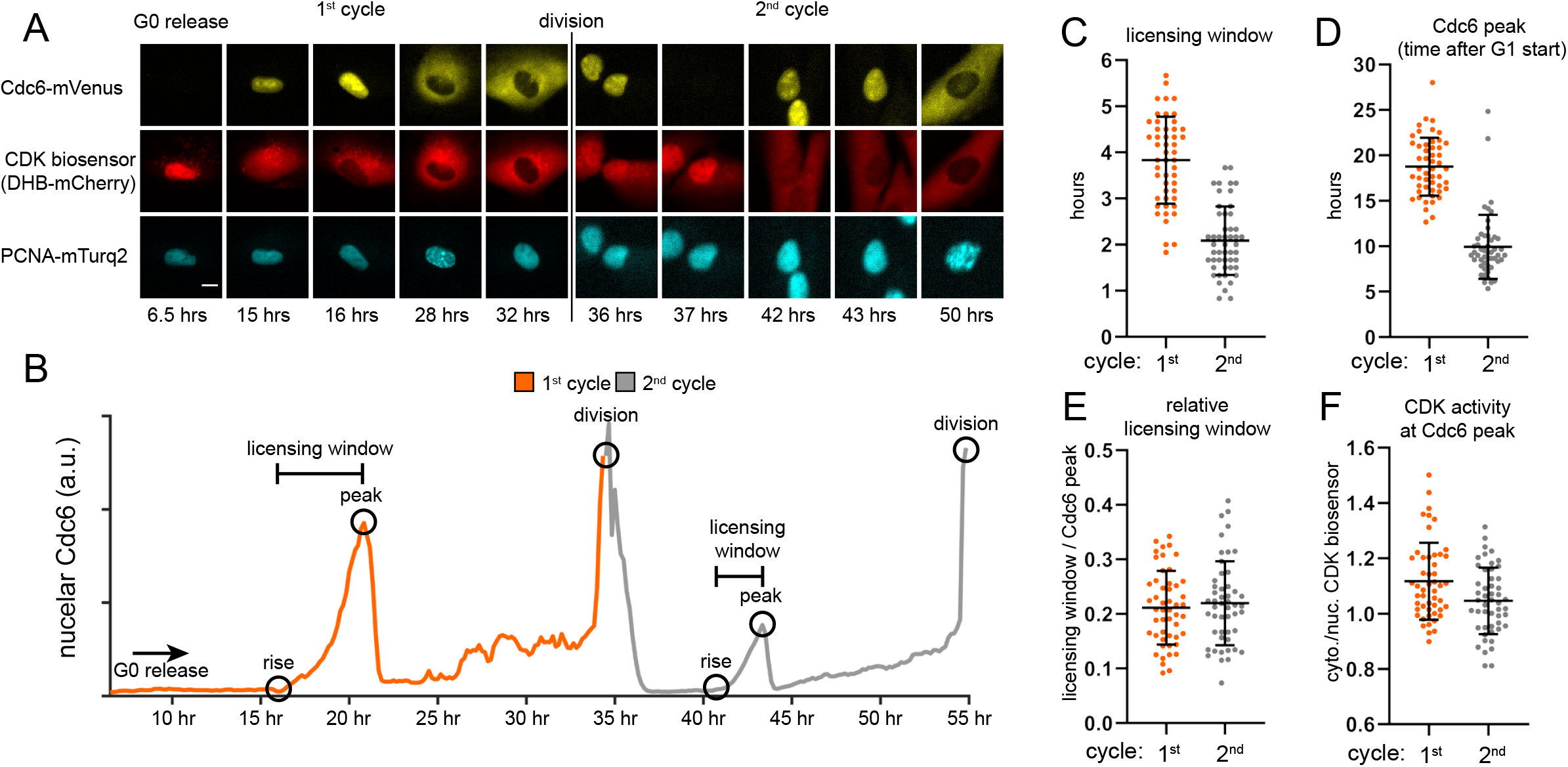
Cdc6 dynamics in the first and second cycles after G0. **a**. RPE1-hTert cells constitutively expressing Cdc6-mVenus, PCNA mTurq2 and a CDK activity sensor (DHB-mCherry), and synchronized in G0 by contact inhibition and release were imaged every 10 minutes for 72 hours. Representative micrographs are shown marking the time of Cdc6 accumulation, CDK activity increase and S phase entry. Images were brightness/contrast adjusted. Representative cell from n=50 analyzed cells. The scale bar is 10 μm and applies to all images. **b**. An individual cell trace of mean nuclear Cdc6 intensity imaged from Fig 6A. The trace is orange for the first cell cycle and grey for one daughter in the cell second cell cycle. Hours indicate hours from G0 release beginning at 6.5 hours after release. Circles indicate relevant features: rise is first appearance of nuclear Cdc6; peak is maximum nuclear Cdc6 before S phase and cytoplasmic translocation. The “licensing window” is the difference between peak and rise. Division is the last image before cytokinesis. **c**. Quantification of licensing window time (Cdc6 peak time minus Cdc6 rise time) for the 50 cells imaged in Fig. 6A, two complete cell cycles for each cell were analyzed. **d**. Timing of Cdc6 peak relative to G0 release (first cell cycle) or to division (second cell cycle) for the 50 cells imaged in Fig. 6A, two cell cycles for each cell. **e**. Ratio of licensing window time divided by Cdc6 peak time for each cell imaged as in Fig. 6A, 50 cells total with both first and second cell cycle **f**. Ratio of mean cytoplasmic DHB-mCherry divided by mean nuclear DHB-mCherry at the time of Cdc6 peak for the 50 cells imaged in Fig. 6A, two cell cycles for each cell.

We calculated the licensing window as a fraction of the total G1 length by dividing the licensing window time by the Cdc6 peak time and found that this window is in fact the same proportion of G1 in both the first and second cell cycles (Fig. 6E). These data suggest that G1 during cell cycle re-entry is a stretched rather than delayed G1 compared to actively proliferating cells in the second cycle, and cells in the first cycle have a longer amount of available time with high Cdc6 than cells in the second cycle. Cells in the first G1 have more potential time to load MCM and yet, enter S phase with less loaded MCM than cells in the second cycle. Taken together, these data suggest that cells in the first G1 after G0 load MCM slowly.

Cells in both the first and second cell cycles also reached a similar CDK activity (cytoplasmic/nuclear ratio greater than 1) at the time of peak nuclear Cdc6 and S phase entry (Fig. 6F) close to previously-reported values of 0.84-1.0 in other untransformed cell lines (Spencer et al., 2013; Schwarz et al., 2018). In other words, the underlicensed first G1 cells achieved the same level of CDK activity in late G1 as normally-licensed second G1 cells. These equivalent CDK activity indicators further support the notion that the licensing checkpoint does not delay CDK2 activation in the first G1.

### Extending the first G1 after G0 substitutes for the impaired licensing checkpoint

Cells re-entering the cell cycle from G0 license origins slowly and begin S phase before they are fully licensed. We hypothesized that artificially extending the first G1 phase could rescue the underlicensing of the first S phase by extending time for MCM loading. If successful, then cells would enter S phase with the high amount of loaded MCM typical of proliferating cells. We chose to extend G1 phase using nutlin-3a, a p53 stabilizing drug previously shown to lengthen G1 (Tovar et al., 2006). We treated cells re-entering the first G1 phase with nutlin-3a beginning in mid-G1 for 8 hours, then washed off the drug to permit passage into S phase and measured the amount of loaded MCM at S phase entry (Fig. 7A, B). Nutlin-3a stabilized p53, causing p21 protein accumulation. This effect is known to inhibit CDK2 kinase activity and delay S phase entry (Giono and Manfredi, 2007). An unwanted secondary effect of low CDK2 activity for our purposes is failure to protect Cdc6 in late G1 phase, which would interfere with our goal of extending the time for MCM loading (Petersen et al., 2000; Mailand and Diffley, 2005). To allow MCM loading during the nutlin-induced G1 arrest, we constitutively expressed a mutant form of Cdc6 that does not require CDK2 activity for stability (Matson et al., 2017). We then compared licensing and cell cycle progression in these treated cells to untreated cells from the first and second cell cycles (Fig. 7C, Fig. S4A). Strikingly, transiently extending the first G1 phase by several hours almost fully rescued licensing in the first cell cycle to the same high level as in the second cell cycle (compare green and grey lines, Fig. 7C, and fold change Fig. 7D) and decreased the percentage of underlicensed cells to the same low level as the second cell cycle (Fig. 7E). Unlike the effects of stable Cdc6 on the rate of MCM loading in proliferating cells (Matson et al., 2017), neither stable Cdc6 alone nor in combination with Cdt1 overproduction increased the rate of MCM loading in the first G1 phase (Fig. S4B and S4C). These genetic changes also had little-to-no effect on underlicensing in the first S phase (Fig. S4D-F). Therefore, the improved licensing by early S phase from nutlin-induced G1 extension is due to the increased time for MCM loading, rather than directly improving MCM loading itself.

**Figure 7.**
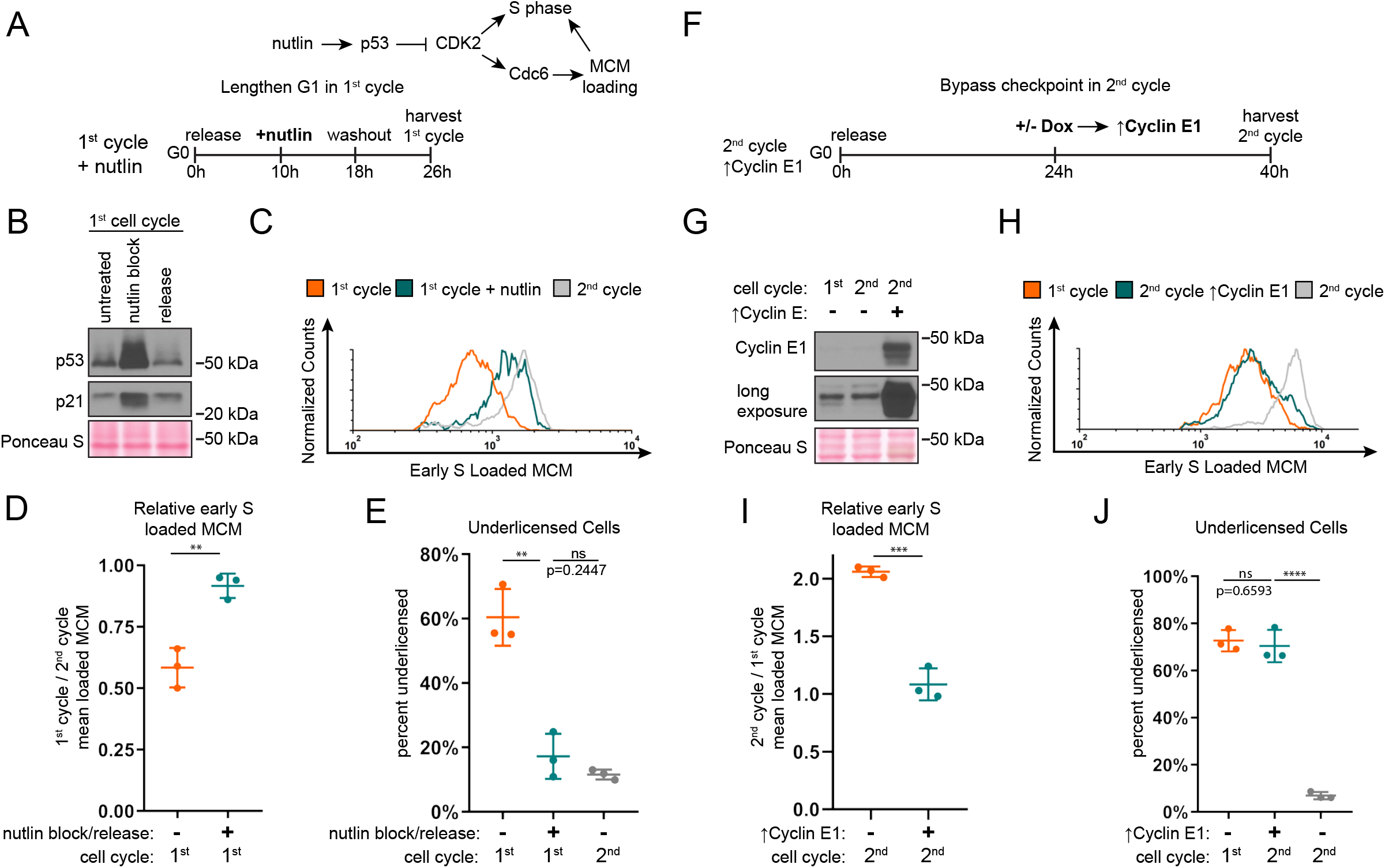
G1 length of first and second cell cycle effects amount of loaded MCM at S phase entry. **a**. Model of experiment. RPE1 cells constitutively producing 5myc-Cdc6-mut (R56A, L59A, K81A, E82A, N83A, not targeted for degradation by APC^CDH1^) were synchronized in G0 by contact inhibition, then released back into the first cell cycle. Cells were treated with 10 uM nutlin-3a 10 hours after release and then nutlin was washed out at 18 hours after release, harvesting cells 26 hours after release from G0. Untreated cells were harvested at 24 hours (first cell cycle) and 48 hours (second cell cycle) after release from G0. **b**. Immunoblot of total protein lysate from cells treated as Fig. 7A. Untreated lane is cells at 18 hours after G0 without nutlin-3a. Nutlin block lane is cells with 10 uM nutlin-3a added at 10 hours and harvested at 18 hours after G0. Release lane is nutlin block, then washed to fresh media and harvested at 26 hours after G0. **c**. Loaded MCM in early S phase determined by flow cytometric analysis from cells treated as Fig. 7A, measuring DNA Content (DAPI), Loaded MCM (anti-Mcm2), and DNA Synthesis (EdU). Histograms show early S loaded MCM. Orange line is first cell cycle, green line is first cell cycle with nutlin block and release, grey line is second cell cycle. See Fig. S4A for complete flow cytometry plots. **d**. Comparison of early S phase DNA-loaded MCM from cells in Fig. 7C. Values plotted are the ratio of mean loaded MCM of first cell cycle with or without nutlin block and release divided by mean loaded MCM of second cell cycle. Horizontal bars indicate means, error bars mark standard deviation (SD), n=3 biological replicates. Samples were compared by unpaired, two tailed t test. p=0.0036**. **e**. Percentage of underlicensed cells from Fig. 7C. Horizontal bars indicate means, error bars mark standard deviation (SD), n=3 biological replicates. Untreated and nutlin treated cells were compared by unpaired, two tailed t test. First cell cycle and nutlin block p=0.0027**, Nutlin block and second cell cycle p=0.2447 (ns). **f**. Diagram of experiment. RPE1 cells containing integrated doxycycline inducible exogenous Cyclin E1 were synchronized in G0 by contact inhibition, then released into the cell cycle, adding 100 ng/mL of doxycycline at 24 hours after release to overproduce Cyclin E1 and shorten G1 of the second cell cycle, harvesting cells at 40 hours after release from G0. Untreated cells were harvested at 24 hours (first cell cycle) and 40 hours (second cell cycle) after release from G0. **g**. Immunoblot for Cyclin E1 on total protein lysate from cells treated as Fig. 7F. **h**. Loaded MCM in early S phase determined by flow cytometric analysis of cells treated as Fig. 7F, measuring DNA Content (DAPI), Loaded MCM (anti-Mcm2), and DNA Synthesis (EdU). Orange line is first cell cycle, green line is second cell cycle with ↑Cyclin E1 starting at 24 hours after G0 release, and grey line is second cell cycle. See Fig. S4G. **i**. Comparison of early S phase loaded MCM per cell from Fig. 7F. Values plotted are the ratio mean loaded MCM of second cell cycle divided by mean loaded MCM of first cell cycle. Horizontal bars indicate means, error bars mark standard deviation (SD), n=3 biological replicates. Samples were compared by unpaired, two tailed t test. p=0.0036** **j**. Percentage of underlicensed cells from Fig. 7F. Horizontal bars indicate means, error bars mark standard deviation (SD), n=3 biological replicates. Samples were compared by unpaired, two tailed t test. first cell cycle vs second cell cycle with ↑Cyclin E1 p=0.6593 (ns), second cell cycle with ↑Cyclin E1 vs second cell cycle untreated p<0.001****.

Finally, we predicted that since the second cell cycle has a robust origin licensing checkpoint, bypassing the checkpoint and artificially shortening the second G1 would induce a phenotype similar to the underlicensed first cell cycle. To test this prediction, we overproduced Cyclin E1 as cells approached the second cell cycle to shorten the second G1 phase and bypass the checkpoint. We then compared licensing in the second S phase in these Cyclin E1-overproducing cells to both control second S phase and first S phase cells (Fig. 7F, 7G, and Fig. S4G). The second cell cycle with a bypassed licensing checkpoint strongly resembled the first cell cycle (compare green and orange lines, Fig. 7H, fold change in Fig. 7I, and percentage of underlicensed cells in Fig. 7J). Taken together, we conclude that cells re-entering the cell cycle from G0 are routinely underlicensed, but that the second and subsequent cell cycles are fully licensed because of a robust p53-dependent checkpoint that ultimately controls the timing of Cyclin E/CDK2-mediated S phase entry.

## DISCUSSION

The origin licensing checkpoint protects cells from premature S phase entry that could lead to genome instability (Fig. 8). This checkpoint couples the activity of CDK to the status of MCM loading such that the G1 CDKs are not activated while MCM loading is still low. Cyclin E1 overproduction can bypass the checkpoint-induced delay and induce premature S phase entry while cells are still underlicensed (Fig. 3) (Ekholm-Reed et al., 2004; Matson et al., 2017). Moreover, p53 loss clearly impairs the normal coupling of MCM loading and S phase entry whereas p53 activation lengthens G1 to give enough time to complete licensing (Fig. 4, 5, and 7). These observations fit Hartwell and Weinert’s classic definition for a cell cycle checkpoint (Hartwell and Weinert, 1989). A checkpoint enforces the dependence of one event on a previous event, but the dependency can be bypassed by mutation to occur out of order. The ability of mutations to induce inappropriate cell cycle progression indicates that in wild-type cells, progression <u>could</u> have occurred but was instead restrained by the checkpoint. This relationship is in contrast to a simple inability to progress. Of relevance to the origin licensing checkpoint, we induced S phase entry when the amount of loaded MCM was low, but not completely absent, since RNAi-mediated depletion of MCM loading factors does not prevent <u>all</u> MCM loading. If we could have entirely prevented MCM loading, then of course, there would have been no DNA synthesis even in p53 null or Cyclin E-overproducing cells because MCM is an essential component of the replicative helicase.

**Figure 8.**
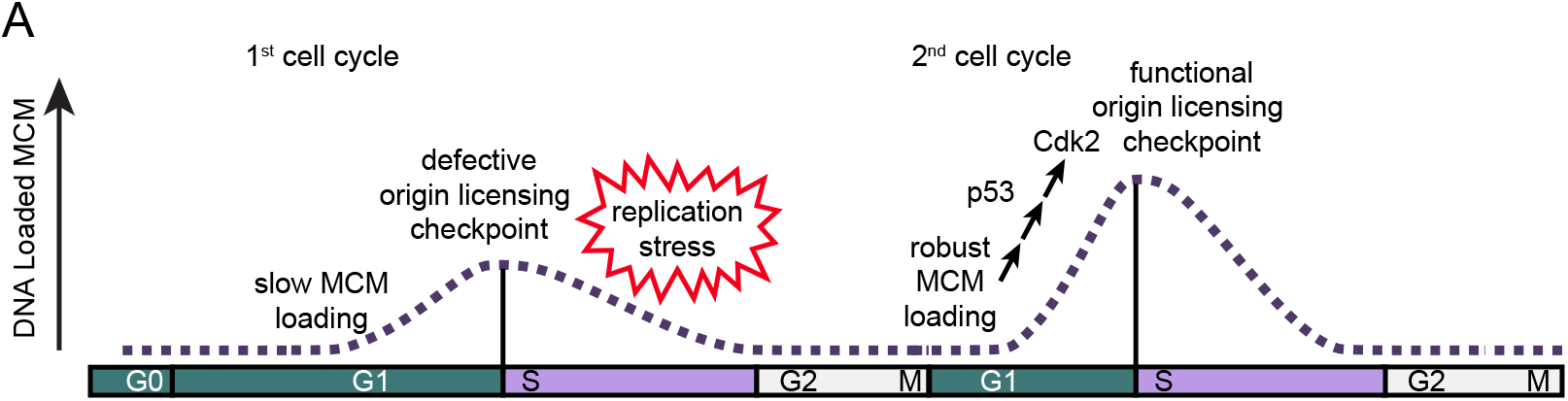
Model of MCM loading in first and second cycles after G0. **a**. Cells in the first cell cycle have slow MCM loading and an impaired origin licensing checkpoint. These defects cause underlicensing and make the first S phase hypersensitive to replication stress. Cells in the second cycle have a functional origin licensing checkpoint and load MCM normally. The checkpoint is p53-dependent and robust MCM loading allows CDK2 activation and normal S phase entry.

Many questions remain about the origin licensing checkpoint itself. How do cells monitor the status of origin licensing, what conditions satisfy the checkpoint, and how is licensing status coupled to the activation of CDKs? Our results indicate that a 50% reduction in the amount of loaded MCM by the time of S phase entry sensitizes human cells to exogenous replication stress; this level is similar to that reported for checkpoint-deficient U2OS cells (Ge et al., 2007). Some stem cell types have a greater amount of loaded MCM at S phase entry than their differentiated counterparts, suggesting that the licensing threshold may vary by cell type (Ge et al., 2015; Carroll et al., 2018). The mere existence of the licensing checkpoint was an unresolved question until relatively recently in large part because it appears to be restricted to untransformed metazoan cells (Piatti et al., 1995). This restriction poses technical challenges for experimentally interrogating the checkpoint because untransformed cells are more difficult to synchronize and manipulate than tumor-derived cells or yeast cells. The fact that p53 status correlates with licensing checkpoint proficiency gives clues that a p53-dependent process, perhaps one or more p53-regulated genes, is involved in efficiently coupling licensing to S phase entry. The discovery here that cell cycle re-entry induces natural underlicensing also suggests future avenues to identify checkpoint mediators. Our analysis of origin licensing status at the G1/S transition using analytical flow cytometry provides a promising opportunity to dissect this pathway in the future.

We note that licensing checkpoint activity is only readily detected in cells with particularly slow MCM loading. In previous studies, MCM loading was artificially slowed to test for checkpoint proficiency in various genetic backgrounds (Shreeram et al., 2002; Liu et al., 2009; Nevis et al., 2009). Under normal growth conditions however both checkpoint-proficient and checkpoint-impaired cell lines enter S phase with normal high levels of MCM loading. This behavior suggests that MCM loading is usually fast enough to achieve high levels of loaded MCM in advance of the other rate-limiting molecular events necessary to trigger CDK2 activation and S phase entry. It is only slow-loading cells that are visibly affected by the checkpoint. In that sense, the origin licensing checkpoint resembles other cell cycle checkpoints that operate in all cell cycles in that “…elimination of the checkpoint may have catastrophic or subtle consequences depending on the prevailing conditions.”(Hartwell and Weinert, 1989) For example, the contribution of the mitotic spindle assembly checkpoint is most obvious when attachments are severely compromised (Lara-Gonzalez et al., 2012). The effect of the spindle assembly checkpoint on the rate of unperturbed mitotic progression is relatively minor except in individual cells that fail to form kinetochore-microtubule attachments on time (Meraldi et al., 2004). Similarly, we did detect a modest increase in the number of underlicensed S phase cells in otherwise unperturbed proliferating p53 null cells compared to p53 WT cells (Fig. 4). We infer that these individual cells loaded MCM slowly, but then could not effectively delay S phase entry.

We were surprised to find that the first G1 phase after cell cycle re-entry from quiescence (G0) is routinely followed by an underlicensed S phase in multiple cell lines. To our knowledge, cell cycle re-entry from G0 is the first known naturally-occurring underlicensed cell cycle; previous investigations required siRNA or mutations to artificially force underlicensing (McIntosh and Blow, 2012). Clearly the extra time in G1 after G0 was not enough to lead to full licensing. Our investigation leads us to conclude that the routine underlicensing in cells re-entering the cycle is the consequence of a combination of slow MCM loading plus an ineffective licensing checkpoint. If MCM loading were fast enough in the first cell cycle, then their checkpoint deficiency wouldn’t matter and they would not be underlicensed in the first S phase. Although it is not yet technically possible to monitor MCM loading in real time in individual cells, we used the transition from Cdc6-negative to Cdc6-positive as a proxy for the maximum amount of time available for MCM loading (Fig. 6). Since cells in the first G1 phase had <u>more</u> time with abundant nuclear Cdc6 than cells in the second G1 phase, yet began S phase with <u>less</u> MCM loaded, we are confident in our assertion that MCM loading is particularly slow in the first G1 phase.

It is unclear why cells re-entering G0 load MCM slowly since it isn’t simply the abundance of Cdc6. The difference may be caused by one or more of the fundamental differences in G1 regulation between cell cycle re-entry and active proliferation. These differences include the status of Rb phosphorylation which is monophosphorylated in proliferating G1 cells, but unphosphorylated in G0 (Narasimha, Kaulich, Shapiro et al., 2014). Rb-dependent genes include those whose products are directly involved in MCM loading, and these genes are subject to unique transcriptional repression by the DREAM complex in G0 that must be reversed at cell cycle re-entry (Litovchick et al., 2007). Proliferating cells also load MCM both before and after Rb is removed from chromatin, but cells re-entering G1 from G0 only load MCM after Rb is removed from chromatin (Håland et al., 2015). CDK2 is constitutively nuclear in proliferating cells, but is cytoplasmic in G0 (Brown et al., 2004; Dietrich et al., 1997). Of additional particular relevance is the indication of a kinase-independent function of Cyclin E1 to promote loading that is only required in cells re-entering the cycle form G0 but not in other cell cycles (Geng et al., 2003, 2007). These differences illustrate that cell cycle re-entry into G1 from G0 poses additional challenges and requirements not encountered during G1 in actively proliferating cells. It is striking however that both the checkpoint defect and slow MCM loading are normal just one cell cycle later after mitosis.

Previous analysis by Daigh et al. indicated that cells released from G0 had increased endogenous replication stress in the first S phase compared to actively proliferating cells. This stress was detected as mid-S phase fluctuations in CDK activity plus markers of the replication stress response signaling pathway (Daigh et al., 2018). Our findings here suggests a molecular mechanism by which that replication stress is generated. We propose that the underlicensed S phase after the first G1 has higher endogenous replication stress because fewer dormant origins are available. While we did not detect frank DNA damage markers in the absence of exogenous replication stress in our cells, we presume that endogenous stress was generated because these cells were uniquely sensitive to low dose gemcitabine. This hypersensitivity is a hallmark of cells that enter S phase while underlicensed (Woodward et al., 2006; Zimmerman et al., 2013). The increased endogenous replication stress from too few licensed origins promotes genome instability from a higher frequency of stalled and unrescued replication forks.

How frequently do cells experience this naturally-underlicensed cell cycle *in vivo?* Cell cycle re-entry from quiescence is common in tissues that naturally turn over cell populations (Sosa et al., 2014; Sagot and Laporte, 2019; Wells et al., 2013). One might imagine however, that one risky cell cycle is a small contributor to overall genome instability. On the other hand, even in actively proliferating cultures, recent studies have detected populations of cells that appear to spontaneously and transiently exit to a quiescence-like state (Spencer et al., 2013; Overton et al., 2014; Arora et al., 2017; Barr et al., 2017). Such transient cell cycle exit may be even more common in tissues compared than in cell culture conditions that have been optimized for maximal uninterrupted growth. Whether or not that transient quiescence also causes underlicensing is not yet known.

Quiescent hematopoietic stem cells re-enter the cell cycle in response to stresses such as viral infection and then return to quiescence (Cheung and Rando, 2013). Interestingly, hematopoietic stem cells acquire DNA damage in the first cycle after G0, accumulate DNA damage over time in older stem cells, and become depleted as they age from repeated cell cycle re-entry (Beerman et al., 2014; Walter et al., 2015). Additionally, quiescent human T cells re-enter the first S phase even when treated with siRNA to substantially reduce the amount of loaded MCM to ~5-10% of their normal loaded amount (Orr et al., 2010). Taken together, these data suggest that at least some quiescent cells *in vivo* lack a licensing checkpoint in the first cycle, although it is unknown if the first cell cycle is naturally underlicensed compared to the second cycle *in vivo*. Old hematopoietic stem cells experience more replication stress in the first S phase after G0 compared to young stem cells, possibly because they express less MCM protein. In that regard, a defective licensing checkpoint would be particularly toxic to old quiescent cells re-entering G1 because they may enter S phase with even less loaded MCM than young cells (Flach et al., 2014). These *in vivo* studies are consistent with our observation that artificially inducing repeated rounds of quiescence and cell cycle re-entry enhanced replication stress sensitivity (Fig. S2). We suggest that over many rounds of quiescence and re-entry, incremental DNA damage accrues from unresolved replication stress or incomplete replication that may pass through mitosis into the next cell generation. The ability of unresolved replication stress to carry forward into subsequent cell cycles was recently established both by artificially inducing underlicensing and by monitoring spontaneous replication stress passing from mother to daughter cells (Moreno et al., 2016; Yang et al., 2017; Arora et al., 2017; Ahuja et al., 2016). Repeated rounds of underlicensed cell cycle re-entry could contribute to the genome damage that drives both aging and oncogenesis. Such accumulating damage could be analogous to the enhanced wear on an automobile that is routinely driven under stop-and-start conditions vs one driven at steady speeds. By that analogy, the most important consequences of entering a high-risk underlicensed cell cycle may appear not in a single cycle, but rather over an organism’s lifetime.

## MATERIALS AND METHODS

### Cell culture and synchronization

RPE1-hTERT cells, 293T and NHF1-hTERT cells (ATCC) were grown in Dulbecco’s modified Eagle’s medium (DMEM) (Sigma Aldrich) with 10% Fetal Bovine Serum (FBS) (Seradigm) and 2 mM L-glutamine (Gibco) at 37°C with 5% CO_2_. Wi38 cells (ATCC) were grown in Minimum Essential Medium (MEM) with 10% FBS (Seradigm), 2 mM L-glutamine (Gibco) and Minimum Essential Medium non-essential amino acids (NEAA) (Gibco) at 37°C with 5% CO_2_. Cells were passaged every three days with trypsin (Sigma Aldrich) and not allowed to reach confluency.

To synchronize RPE1 cells in G0 by contact inhibition, cells were grown to 100% confluency, washed with PBS, and incubated in DMEM with 10% FBS and 2 mM L-glutamine for 48 hours. To release cells into the first or second cell cycles, G0 cells were re-stimulated by passaging 1:10 for the first cycle, harvesting 24 hours later or 1:20 for the second cycle, harvesting 48 hours later, in DMEM with 10% FBS and 2 mM L-glutamine. To synchronize RPE1 cells in G0 by serum starvation, cells were plated sub-confluent, washed 3 times with PBS, and starved in DMEM with 0% FBS and 2 mM L-glutamine for 72 hours. To release cells into the first or second cell cycles, cells were washed with PBS and re-stimulated by washing once with PBS and adding DMEM with 10% FBS and 2mM L-glutamine for 24 hours (first cycle) or 48 hours (second cycle). To synchronize NHF1-hTERT cells in G0, cells were grown to 100% confluency, washed with PBS, and incubated in DMEM with 0.1% FBS and 2 mM L-glutamine for 72 hours. To release cells into the first or second cell cycles, G0 cells were re-stimulated by passaging 1:4 for first cycle, harvesting 24 hours later or 1:8 for second cycle, harvesting 48 hours later, in DMEM with 10% FBS and 2 mM L-glutamine. To synchronize Wi38 cells in G0, cells were grown to 100% confluency, washed with PBS, and incubated in MEM with 0.1% FBS, 2 mM L-glutamine, and NEAA for 72 hours. To release cells into the first or second cell cycles, G0 cells were re-stimulated by passaging 1:4 for first cycle, harvesting 24 hours later or 1:8 for second cycle, harvesting 48 hours later, in in MEM with 10% FBS, 2 mM L-glutamine, and NEAA.

To synchronize RPE1 in 3 repeated G0s, cells were synchronized in G0 by contact inhibition (above), re-stimulated into the cell cycle by passaging 1:6 in DMEM with 10% FBS and 2 mM L-glutamine, and grown to 100% confluency within 48-72 hours to start the second G0 by contact inhibition for 48 hours. Cells were re-stimulated as before and grown to 100% confluency to start the 3^rd^ G0 for 48 hours. To release cells for the experiment, G0 cells were re-stimulated by passaging 1:10 in DMEM with 10% FBS and 2 mM L-glutamine, harvesting the first cell cycle at 24 hours after restimulation.

### DNA Cloning and cell lines

The RPE1 CRISPR p53 KO was a gift of Dr. Prasad Jallepalli and described previously (Rodriguez-Rodriguez et al., 2018). The RPE1 cells with doxycycline inducible Cyclin E1 and the RPE1 cells with doxycycline inducible Cdt1 and stable Cdc6 WT or Cdc6-mut were described previously (Matson et al., 2017). The pLenti-PGK hygro PCNA-mTurq2 was described previously (Grant et al., 2018). The CSII-EF zeo DHB-mCherry CDK activity reporter was a gift Dr. Sabrina Spencer. The CSII-EF zeo Cdc6-mVenus was a gift from Dr. Michael Brandeis. Each reporter is under control of a constitutive heterologous promoter.

To make the RPE1 line containing PCNA-mTurq2, Cdc6-mVenus, and DHB-mCherry, the plasmids were transfected into 293T with pCI-GPZ or ΔNRF and VSVG virus packaging plasmids with 50 ug/mL Polyethylenimine (Aldrich Chemistry). Viral supernatants were transduced onto RPE1 cells with 8 ug/mL Polybrene (Millipore). A clonal cell line was picked based on fluorescence of all 3 biosensors.

### siRNA transfections and drug treatment

siRNA concentration and sequences:

siControl, (siLuciferase) 100 nM (synthesized by Life Technologies)
5’-CUUACGCUGAGUACUUCGA-3’
siCdt1 A, mixture of 4 sequences, 25 nM each. (siGENOME CDT1 siRNA, Dharmacon)
5’-CCAAGGAGGCACAGAAGCA-3’
5’-GCUUCAACGUGGAUGAAGU-3’
5’-UCUCCGGGCCAGAAGAUAA-3’
5’-GGACAUGAUGCGUAGGCGU-3’
siCdt1 B, 50 nM (synthesized by Life Technologies)
5’-CCUACGUCAAGCUGGACAATT-3’

For siRNA transfections, siRNA was spotted into plates with DharmaFECT 4 transfection reagent (Dharmacon) and Opti-MEM (Gibco), incubating for 20 minutes before adding cells in DMEM with 10% FBS and 2 mM L-glutamine (final concentrations), harvesting cells 72 hours later for proliferating experiments, and 24 hours later for G0 release, as described above. To overproduce Cyclin E1 or Cdt1-HA, 100 ng/mL of doxycycline (CalBiochem) was added at times indicated in the figure legend. To block and release cells with nutlin-3a (Sigma Aldrich), RPE1 were synchronized in G0 by contact inhibition, released into the first cell cycle, adding 10 uM nutlin-3a 10 hours after release, washing off nutlin-3a 18 hours after release by washing 3 times with PBS and adding back DMEM with 10% FBS and 2 mM L-glutamine. To treat cells with gemcitabine (Sigma Aldrich) in Fig. 2, cells were synchronized in G0 by contact inhibition, released into the first cycle and treated with 50 nM gemcitabine from 22-24 hours or released into the second cycle and treated with 50 nM gemcitabine from 44-46 hours. To treat cells with gemcitabine in Fig. S2, cells were synchronized in one or three G0 (above), adding 5 nM gemcitabine at time of re-stimulation from 0-24 hours.

### Total protein lysate and chromatin fractionation

To prepare total protein lysate for immunoblot, cells were harvested with trypsin and frozen in dry ice, then lysed in cold cytoskeletal buffer (CSK) (10 mM Pipes pH 7.0, 300 mM sucrose, 100 mM NaCl, 3 mM MgCl_2_ hexahydrate) with 0.5% triton x-100 (Sigma Aldrich) and protease and phosphatase inhibitors (0.1 mM Pefabloc, 1 μg/mL pepstatin A, 1 μg/mL leupeptin, 1 μg/mL aprotinin, 10 μg/mL phosvitin, 1 mM β-glycerol phosphate, 1 mM sodium orthovanadate) on ice for 15 minutes. Lysate was centrifuged at 13,000 × g at 4°C for 10 minutes, and a Bradford Assay (Biorad) was done on the supernatant to load equal amounts of protein per sample.

To prepare chromatin fractions for immunoblot, cells were harvested with trypsin and frozen in dry ice, then lysed in cold CSK with 0.5% triton x-100, 1 mM ATP, 5 mM CaCl_2_ and protease and phosphatase inhibitors (complete CSK) on ice for 20 minutes. Then a Bradford assay for equal loading was done on the lysate, a small aliquot removed from each sample as total lysate, and complete CSK was added to each sample, mixed, and centrifuged for 5 min, 1,000 × g. The supernatant was removed, pellet was washed again with complete CSK, incubated for 10 minutes on ice, and then centrifuged for 5 min, 1,000 × g. The supernatant was removed, and DNA loaded proteins were released by incubation with S7 nuclease (Sigma Aldrich) in complete CSK at room temperature (RT) for 10 minutes. Samples were centrifuged again, keeping the supernatant as the chromatin fraction.

### Immunoblotting

Samples were diluted with loading buffer to final concentration: 1% SDS, 2.5% 2-mercaptoethanol, 0.1% bromophenol blue, 50 mM Tris pH 6.8, 10% glycerol then boiled. Samples were run on SDS-PAGE gels, then transferred to nitrocellulose (GE Healthcare) or polyvinylidene difluoride membranes (Thermo Fisher). After transferring, samples were blocked in 5% milk in tris buffered saline with 0.1% tween 20 (TBST), then incubated overnight at 4°C in primary antibody with 2.5% milk in TBST. Then membranes were washed with TBST, incubated in horseradish peroxidase conjugated (HRP) secondary antibody for 1 hour at RT, washed with TBST, and imaged with ECL Prime (Amersham) on autoradiography film (Denville). Antibodies were: Mcm2 (BD Biosciences Cat#610700, 1:10,000), Cyclin E1 (Cell Signaling Technology #4129S, 1:2,000), p53 (Santa Cruz Biotechnology, sc-126, 1:2,000), p21 (Cell Signaling Technology, #2947S, 1:6,000) Cdc6 (Santa Cruz Biotechnology, sc-9964, 1:2,000), Cdt1 (Santa Cruz Biotechnology, sc-365305, 1:3,000), Donkey-anti-Mouse-HRP (Jackson ImmunoResearch, 1:10,000) Donkey-anti-Rabbit-HRP (Jackson ImmunoResearch, 1:10,000). Membranes were stained with Ponceau S (Sigma Aldrich) to determine equal protein loading.

### Flow cytometry

Cells stained with EdU were incubated with 10 μM EdU for 30 minutes before harvesting, except in Fig S1D, when cells were incubated with 1 μM EdU from restimulation to harvesting. Cells were harvested with trypsin to measure DNA loaded proteins by flow cytometry. Cells were washed once with PBS, centrifuged at 2,000 × g for 3 minutes, then supernatant was aspirated and pellets lysed on ice in cold CSK with 0.5% triton X-100 and protease and phosphatase inhibitors for 5 minutes. After incubation, 1% Bovine serum albumin (Fisher) in PBS (B-PBS) was added to each sample, mixed, and centrifuged at 2,000 × g for 3 minutes. Supernatant was aspirated, pellets were fixed with 4% paraformaldehyde (Electron Microscopy Sciences) in PBS for 15 minutes at RT. Then B-PBS was added, mixed, and samples centrifuged (2,000 × g for 7 minutes, all following steps). Supernatant was aspirated, B-PBS was added, and samples stored at 4°C before labeling.

EdU labeling was done before antibody staining. Cells were centrifuged, supernatant aspirated, and incubated in PBS with 1 mM CuSO_4_, 100 mM ascorbic acid (fresh), 1 uM Alexa Flour-647-azide (Life Technologies) for 30 minutes, RT in the dark. Then B-PBS with 0.5% NP-40 (United States Biochemical) was added, mixed, and centrifuged. For antibody staining, supernatant was aspirated, cells incubated in primary antibody: anti-Mcm2 (1:200, BD Biosciences, Cat#610700), anti-γH2AX (1:200, Cell Signaling Technologies, #9718S) in B-PBS with 0.5% NP-40 for 1 hour at 37°C in the dark. Next, B-PBS with 0.5% NP-40 was added, mixed, and samples centrifuged. Supernatant was aspirated, cells incubated in secondary antibody: Donkey-anti-Mouse-Alexa Flour 488 (1:1,000, Jackson ImmunoResearch, for Mcm2), Donkey-anti-Rabbit-Alexa Flour 647 (1:1,000, Jackson ImmunoResearch, for γH2AX) in B-PBS with 0.5% NP-40 for 1 hour at 37°C in the dark. Then B-PBS with 0.5% NP-40 was added, mixed, and samples centrifuged. Finally, supernatant was aspirated, cells were incubated in 1 ug/mL DAPI (Sigma Aldrich) and 100 ng/mL RNAse A (Sigma Aldrich) in B-PBS with 0.5% NP-40 overnight at 4°C in the dark. For negative control samples used to draw positive/negative gates, cells were not incubated with EdU but were labeled with Alexa Flour-647-azide, and were not incubated with primary antibody but were labeled with secondary antibody and DAPI.

Data were acquired primarily on an Attune NxT flow cytometer (Thermo Fisher), with some data acquired on a CyAn ADP flow cytometer (Beckman Coulter). Data was analyzed using FCS Express 6 (De Novo Software). Gates are shown in Fig. S1. Gate to isolate cells from debris was from Forward Scatter-Area vs Side Scatter Area. Gate to isolate singlets from doublets was from DAPI Area vs DAPI height (Parent gate: cells). Gate to isolate cell cycle phases was from DAPI Area (DNA Content) vs 647-Area (DNA Synthesis, EdU) (Parent gate: singlets). Color gates to isolate S-MCM^DNA^ positive, G1-MCM^DNA^ positive and MCM^DNA^ negative were on 647 Area (DNA Synthesis, EdU) vs 488 Area (Loaded MCM), using a negative control sample to mark positive cells (Parent gate: singlets). Early S phase gate was on DAPI Area (DNA Content) vs 488-Area (Loaded MCM), gating cells with 2C DNA content, S-MCM^DNA^ positive in early S phase (Parent gate: S-MCM^DNA^ positive). Mid S phase gate was on DAPI Area (DNA Content) vs 647-Area (γH2AX), cells between 2C and 4C DNA Content (Parent gate: singlets). Replication stress induced γH2AX was gated as cells equal to or greater than the top 5%-6% of γH2AX signal from untreated cells (Parent gate: Mid S). Each flow cytometry plot typically has 9,000-11,000 total single cells. Histogram counts were normalized to the peak value of the 2^nd^ cell cycle or siControl. The normalization allows visual comparison of cell distributions between populations with different numbers of cells due to changes in synchrony in the second cell cycle. The quantification of relative loaded MCM or underlicensed cells were not normalized.

### Doubling time

Cells were plated with siRNA and counted 48 hours or 72 hours later after dissociating with trypsin using a Luna II automated cell counter (Logos Biosystems). Each treatment was done with 3 biological replicates and each dish in the replicate was counted twice as technical replicates. Doubling time was calculated using Prism 8 (GraphPad) regression analysis: exponential growth equation. Doubling times from the 3 biological replicates were averaged, and then the average doubling time was multiplied by the cell cycle phase percentages to obtain cell cycle phase hours.

### Live cell imaging

Cells were plated for live cell imaging with G0 by contact inhibition and restimulation in Flourobrite DMEM (Gibco) with 10% FBS and 2 mM L-glutamine in #1.5 glass bottom plates (Cellvis) in a humidified enclosure at 37° C with 5% CO_2_. Movie started 6.5 hours after plating, cells were imaged for 72 hours with images every 10 minutes. Cells were imaged on a Nikon Ti Eclipse inverted microscope with a 20x (NA 0.75) Apochromat dry objective lens with the Nikon Perfect Focus System. The camera was an Ando Zyla 4.2 sCMOS detector with 12 bit resolution, filters were from Chroma: (excitation; beam splitter; emission filter) CFP - 436/20 nm; 455 nm; 480/40 nm, YFP - 500/20 nm; 515 nm; 535/30 nm; and mCherry - 560/40 nm; 585 nm; 630/75 nm. Images were collected with Nikon NIS-Elements AR software.

Images were analyzed with Fiji version 1.51n, ImageJ. Briefly, images were background subtracted and tracked with custom ImageJ scrips based on the PCNA signal, and nuclear signal quantified in a region of interest as described previously (Grant, Kedziora et al., 2018). Cdc6 traces of mean nuclear intensity were scored as follows: Cdc6 peak time: frame with highest nuclear Cdc6 intensity before the sharp drop in intensity indicating export to the cytoplasm. Cdc6 rise time: frame with Cdc6 nuclear intensity 2 standard deviations greater than the lowest Cdc6 nuclear intensity before Cdc6 peak time. Licensing window time: Cdc6 peak time minus Cdc6 rise time. Relative licensing window time: Licensing window time divided by Cdc6 peak time.

CDK biosensor (DHB-mCherry) cytoplasmic measurement was described previously (Chao et al., 2019). Fluorescence signal of the cytoplasm was approximated by measuring signal within a ring-shaped region (5 px wide) around the nucleus. The ratio between the cytoplasm and the nucleus was calculated using mean signals of the ring-shaped cytoplasmic region and the full nuclear region.

## Supporting information

Matson Supp Figures

## SUPPLEMENTAL MATERIAL

The supplemental figures all support the main figures. Figure S1 defines flow cytometry gating and raw data for additional cell lines summarized in Figure 1G and H. Figure S2 demonstrates replication stress sensitivity over repeated rounds of G0 and cell cycle reentry. Figure S3 is complete flow cytometry color dot plots and histograms of G1 MCM loading with siCdt1 treatment. Figure S4 is complete flow cytometry color dot plots for the nutlin-3a block and release as well as Cyclin E1 overproduction in the second cell cycle. It also includes the minimal effects of Cdc6 and Cdt1 overproduction in the first cell cycle on underlicensing.

## ACKNOWLEDGEMENTS

We thank Jeffrey Jones for laboratory managerial assistance, and members of the Cook lab for helpful discussion of the manuscript. We thank Dr. Sam Wolff, Seraphina Wong, Aimee Littlejohn, and Walli Driggers for lab support. We thank Dr. Kasia Kedziora for help with live cell imaging analysis. The DHB-mCherry plasmid was a generous gift from Dr. Sabrina Spencer. The Cdc6-mVenus plasmid was a generous gift from Dr. Michael Brandeis. The RPE1 p53 CRIPSR null line was a generous gift of Dr. Prasad Jallepalli. This work was supported by a fellowship from the National Science Foundation (DGE-1144081) to J.P.M, a UNC Dissertation Completion Fellowship to J.P.M, grants from the National Institutes of Health/National Institute for General Medical Sciences to J.G.C. (R25GM089569, GM083024, and GM102413). The UNC Flow Cytometry Core Facility is supported in part by P30 CA016086.

The authors declare no competing financial interests.

## AUTHOR CONTRIBUITIONS

J.P.M. designed and performed the experiments and analyzed the results. A.M.H. tracked cells from live cell imaging and processed data. G.D.G. tracked cells and ran the microscope. H.W. performed some immunoblots. J.B.P. created the RPE1 Cdc6-mVenus, PCNA-mTurq2, DHB-mCherry cell line. J.G.C. designed experiments and supervised the project. J.P.M and J.G.C. wrote the manuscript with input from the other authors.

---

**Supplemental Figure 1. Flow cytometry gating and alternate cell lines**

**a**. Flow cytometry gating to isolate cells for analysis. Proliferating RPE1 cells (Fig. 1A) were processed for analytical flow cytometry for chromatin-bound proteins, labeling DNA Content (DAPI), Loaded MCM (anti-Mcm2) and DNA Synthesis (EdU). Cells were labeled with 10 uM EdU for 30 minutes before harvesting. Gating to isolate cells from debris is Forward Scatter Area vs Side Scatter Area, cells gate. Gating to isolate single cells from doublets is DAPI Area vs DAPI Height, singlets gate. Gating to determine cell cycle phase distributions is DNA Content vs DNA Synthesis. Color gating for S-MCM^DNA^ positive (orange), G1-MCM^DNA^ positive (blue) and MCM^DNA^ negative (grey) is on DNA Synthesis vs Loaded MCM using a negative control sample without Mcm2 primary antibody or EdU, but with Donkey anti Mouse-488 secondary antibody and 647-azide as a measure of background staining.

**b**. Cell cycle phase of RPE1 cells G0 synchronized by contact inhibition, Fig. 1C, RPE1 cells G0 synchronized by mitogen starvation, Fig. S1I, Wi38 cells G0 synchronized by contact inhibition, Fig. S1G, and NHF1-htert cells G0 synchronized by contact inhibition, Fig. S1E. Horizontal bars indicate means, error bars mark standard deviation (SD), n=3 biological replicates.

**c**. Immunoblot for Cyclin D1 or p27 on total protein lysate from RPE1 cells synchronized in G0 or released into the first cell cycle (24 hours) as in Fig. 1C.

**d**. Percentage of S phase cells defined by analytical flow cytometry. RPE1 cells were synchronized in G0 by contact inhibition were released into the cell cycle with 1 uM EdU at time of release, harvesting cells, 24, 28, and 32 hours after release from G0. S phase was determined by DAPI (DNA Content) and EdU (DNA Synthesis) cells as in Fig. S1A. Horizontal bars indicate means, error bars mark standard deviation (SD), n=3 biological replicates.

**e**. Flow cytometry of chromatin-bound protein on Wi38 cells synchronized in G0 by contact inhibition in 0.1% FBS for 72 hours and released from G0 into the cell cycle, harvesting 24 hours after release (first cell cycle) and 48 hours after release (second cell cycle). Flow cytometry measured DNA Content (DAPI), Loaded MCM (anti-Mcm2) and DNA Synthesis (EdU). Cells were labeled with 10 μM EdU for 30 minutes before harvesting. Orange cells are S phase MCM^DNA^ positive, blue cells are G1 phase MCM^DNA^ positive, grey cells are MCM^DNA^ negative.

**f**. Loaded MCM in early S phase determined by flow cytometric analysis of early S phase Wi38 from Fig. S1E. Orange line is first cell cycle, grey line is second cell cycle.

**g**. Flow cytometry of chromatin-bound protein on NHF1-htert cells synchronized in G0 by contact inhibition in 0.1% FBS for 72 hours and released from G0 into the cell cycle, harvesting 24 hours after release (first cell cycle) and 48 hours after release (second cell cycle). Flow cytometry measured DNA Content (DAPI), Loaded MCM (anti-Mcm2) and DNA Synthesis (EdU). Orange cells are S phase MCM^DNA^ positive, blue cells are G1 phase MCM^DNA^ positive, grey cells are MCM^DNA^ negative.

**h**. Loaded MCM in early S phase determined by flow cytometric analysis of NHF1 from Fig. S1G. Orange line is first cell cycle, grey line is second cell cycle.

**i**. Flow cytometry of chromatin-bound protein on RPE1 cells synchronized in G0 by starvation in 0% FBS for 72 hours and released from G0 into the cell cycle, harvesting 24 hours after release (first cell cycle) and 48 hours after release (second cell cycle). Flow cytometry measured DNA Content (DAPI), Loaded MCM (anti-Mcm2) and DNA Synthesis (EdU). Orange cells are S phase MCM^DNA^ positive, blue cells are G1 phase MCM^DNA^ positive, grey cells are MCM^DNA^ negative.

**j.** Loaded MCM in early S phase determined by flow cytometric analysis of RPE1 from Fig. S1I. Orange line is first cell cycle, grey line is second cell cycle.

**Supplemental Figure 2. Repeated transitions between G0 and proliferation trend towards an increased replication stress sensitivity**

**a**. Flow cytometry of chromatin-bound protein from cells in Fig. 2B, showing DNA loaded MCM (anti-MCM2). Red cells are gemcitabine (replication stress) induced γH2AX positive, as indicated in Fig. 2C.

**b**. Diagram of experiment. For 1xG0: RPE1 cells were synchronized in G0 by contact inhibition for 48 hours, then released into the cell cycle with 5 nM gemcitabine, harvesting cells 24 hours after release. For 3xG0: RPE1 cells were synchronized in G0 by contact inhibition for 48 hours, released into the cell cycle without gemcitabine and grown to confluency to repeat the G0 by contact inhibition a second time, then repeating once again for a 3^rd^ G0. Then cells were released into the cell cycle with 5 nM gemcitabine, harvesting cells 24 hours after release.

**c**. Flow cytometry of chromatin-bound protein from cells treated as in Fig. S2B, measuring DNA Content (DAPI) and γH2AX (anti H2AX phospho S139). Histograms plot γH2AX of Mid-S phase cells as described in Fig 2C. Upper panel is 1xG0, lower panel is 3xG0. Grey lines are first cell cycle untreated, red lines are first cell cycle treated with 5 nM gemcitabine.

**d**. Percentage of replication stress-induced γH2AX in 1xG0 and 3xG0 release into the first cycle from Fig. S2C. Horizontal bars indicate means, error bars mark standard deviation (SD), n=3 biological replicates. 1xG0 and 3xG0 compared by unpaired, two tailed t test. p=0.153 (ns).

**e**. Comparison of γH2AX intensity per cell presented as fold-change between gemcitabine-treated and untreated cells from Fig. S2C. Horizontal bars indicate means, error bars mark standard deviation (SD), n=3 biological replicates. Samples compared by unpaired, two tailed t test. p=0.1464 (ns).

**Supplemental Figure 3. Flow cytometry plots of siRNA in proliferating cells.**

**a**. Flow cytometry of chromatin-bound protein of cells shown in Fig. 3C, measuring DNA Content (DAPI), Loaded MCM (anti-Mcm2) and DNA Synthesis (EdU). Orange cells are S phase MCM^DNA^ positive, blue cells are G1 phase MCM^DNA^ positive, grey cells are MCM^DNA^ negative.

**b**. Loaded MCM in G1 cells from Fig 3C. Black lines are siControl treated cells. Green lines are siCdt1 A (left) or siCdt1 B (right). Blue lines are siCdt1 A, ↑Cyclin E1 (left) or siCdt1 B, ↑Cyclin E1 (right).

**c**. Flow cytometry of chromatin-bound protein of cells shown in Fig. 4C, measuring DNA Content (DAPI), Loaded MCM (anti-Mcm2) and DNA Synthesis (EdU). Orange cells are S phase MCM^DNA^ positive, blue cells are G1 phase MCM^DNA^ positive, grey cells are MCM^DNA^ negative.

**d**. Loaded MCM in G1 cells from Fig. 4C. Left histogram: Black line is siControl, p53 WT, blue line is siControl p53 KO. Middle and right histograms: Black lines are siControl, p53 WT, green lines are siCdt1A, p53 WT (left) or siCdt1 B, p53 WT (right), blue lines are siCdt1 A p53 KO (left) or siCdt1 B p53 KO (right). Note the black siControl p53 WT are the same sample on all three histograms.

**Supplemental Figure 4. Overproduction of Cdt1 and a stable Cdc6 mutant do not rescue underlicensing in the first cell cycle after G0**.

**a**. Flow cytometry of chromatin-bound protein from cells shown in Fig. 7C, measuring DNA Content (DAPI), Loaded MCM (anti-Mcm2) and DNA Synthesis (EdU). 1^st^ cell cycle is 24 hrs after G0 release, 1^st^ cycle with nutlin block and release are cells treated with 10 μM nutlin-3a from 10-18 hours and released to 26 hours after G0. 2^nd^ cell cycle is 48 hours release from G0. Orange cells are S phase MCM^DNA^ positive, blue cells are G1 phase MCM^DNA^ positive, grey cells are MCM^DNA^ negative.

**b**. Immunoblot for Cdc6 and Cdt1 on total protein lysate from RPE1 cells constitutively producing either 5myc-Cdc6 WT or 5myc-Cdc6-mut (a mutant of Cdc6 that is not targeted for degradation by APC^Cdh1^: R56A, L59A, K81A, E82A, N83A) and a doxycycline inducible Cdt1-HA. RPE1 cells were synchronized in G0 by contact inhibition, treated with 100 ng/mL doxycycline for 4 hours before re-plating cells to release into the first cell cycle, harvesting cells 24 hours after release.

**c**. Flow cytometry of chromatin-bound protein on cells treated as Fig. S4B, harvested 24 hours (first cell cycle) and 48 hours (second cell cycle) after G0 release, measuring DNA Content (DAPI), Loaded MCM (anti-Mcm2), and DNA Synthesis (EdU). Orange cells are S phase MCM^DNA^ positive, blue cells are G1 phase MCM^DNA^ positive, grey cells are MCM^DNA^ negative.

**d**. Loaded MCM of early S phase from Fig. S4C. Orange lines are cells from first cell cycle, grey lines are cells from second cell cyclist. Top panel is RPE1 cells producing Cdt1-HA WT and 5myc-Cdc6 WT, bottom panel is RPE1 cells producing Cdt1-HA WT, 5myc-Cdc6-mut.

**e**. Comparison of early S phase DNA-loaded MCM per cell from Fig. S4D. Values plotted are the ratio of mean loaded MCM of first cell cycle divided by mean loaded MCM of second cell cycle. Horizontal bars indicate means, error bars mark standard deviation (SD), n=3 biological replicates. First and second cell cycles compared by unpaired, two tailed t test. p=0.8635 (ns).

**f**. Percentage of underlicensed cells from early S phase cells in Fig. S4D. Horizontal bars indicate means, error bars mark standard deviation (SD), n=3 biological replicates. First and second cell cycles compared by unpaired, two tailed t test. Cdc6 WT p=0.0041** Cdc6 mut p=0.0001***.

**g**. Flow cytometry of chromatin-bound protein from cells shown in Fig. 7H, measuring DNA Content (DAPI), Loaded MCM (anti-Mcm2) and DNA Synthesis (EdU). Orange cells are S phase MCM^DNA^ positive, blue cells are G1 phase MCM^DNA^ positive, grey cells are MCM^DNA^ negative.

## Abbreviations

Cdc6: Cell division cycle 6
CDK: Cyclin dependent kinase
Cdt1: Cdc10 dependent transcript 1
KO: knockout
MCM: Minichromosome maintenance complex

