## Supplementary material for "Intrinsic checkpoint deficiency during cell cycle re-entry from quiescence": Matson Supp Figures

Figure S1, Matson et al.

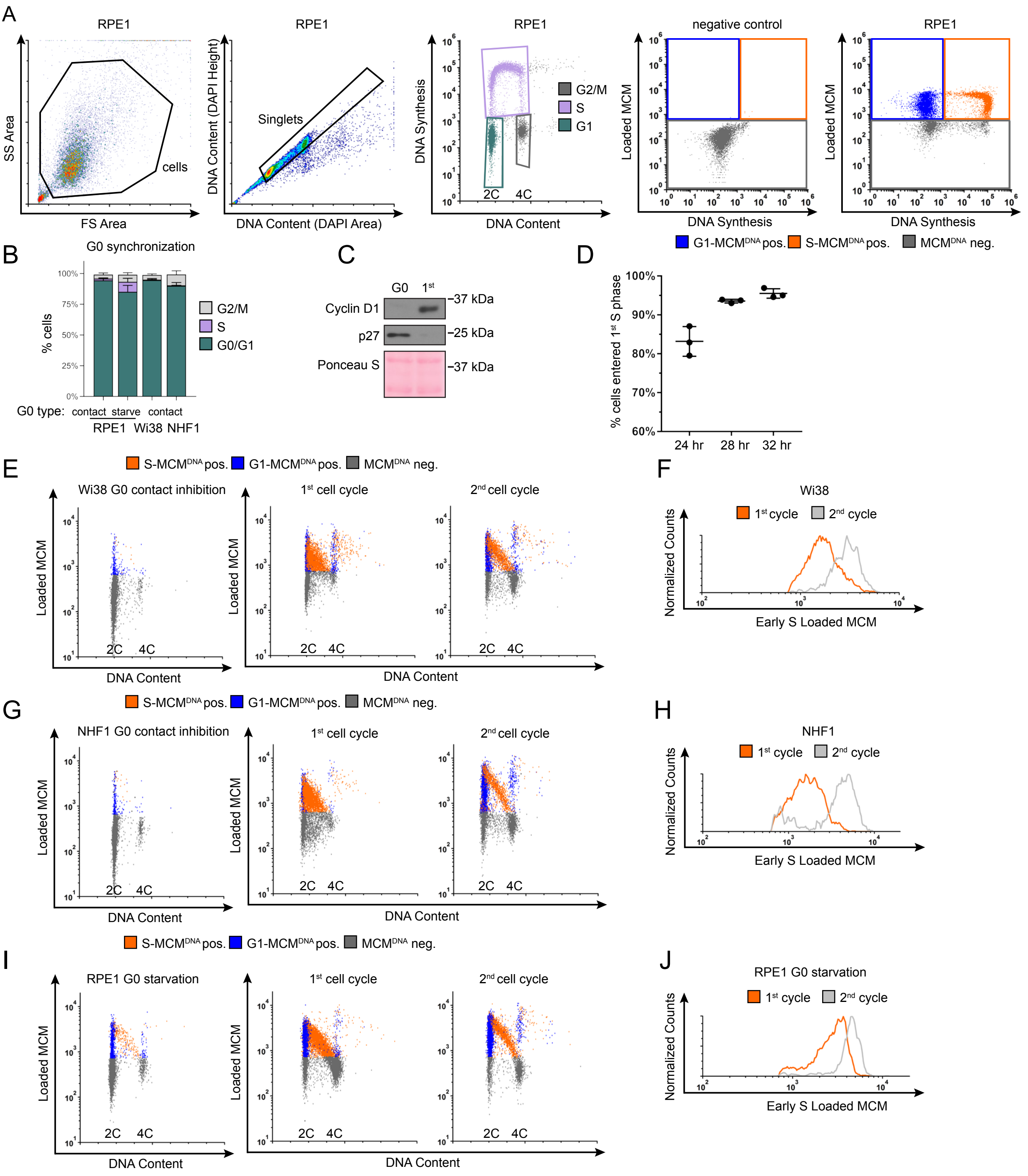

**Supplemental Figure 1. Flow cytometry gating and alternate cell lines**
**a.** Flow cytometry gating to isolate cells for analysis. Proliferating RPE1 cells (Fig. 1A) were processed for analytical flow cytometry for chromatin-bound proteins, labeling DNA Content (DAPI), Loaded MCM (anti-Mcm2) and DNA Synthesis (EdU). Cells were labeled with 10 uM EdU for 30 minutes before harvesting. Gating to isolate cells from debris is Forward Scatter Area vs Side Scatter Area, cells gate. Gating to isolate single cells from doublets is DAPI Area vs DAPI Height, singlets gate. Gating to determine cell cycle phase distributions is DNA Content vs DNA Synthesis. Color gating for S-$MCM^{DNA}$ positive (orange), G1-$MCM^{DNA}$ positive (blue) and $MCM^{DNA}$ negative (grey) is on DNA Synthesis vs Loaded MCM using a negative control sample without Mcm2 primary antibody or EdU, but with Donkey anti Mouse-488 secondary antibody and 647-azide as a measure of background staining.
**b.** Cell cycle phase of RPE1 cells G0 synchronized by contact inhibition, Fig. 1C, RPE1 cells G0 synchronized by mitogen starvation, Fig. S1I, Wi38 cells G0 synchronized by contact inhibition, Fig. S1G, and NHF1-htert cells G0 synchronized by contact inhibition, Fig. S1E. Horizontal bars indicate means, error bars mark standard deviation (SD), n=3 biological replicates.
**c.** Immunoblot for Cyclin D1 or p27 on total protein lysate from RPE1 cells synchronized in G0 or released into the first cell cycle (24 hours) as in Fig. 1C.
**d.** Percentage of S phase cells defined by analytical flow cytometry. RPE1 cells were synchronized in G0 by contact inhibition were released into the cell cycle with 1 uM EdU at time of release, harvesting cells, 24, 28, and 32 hours after release from G0. S phase was determined by DAPI (DNA Content) and EdU (DNA Synthesis) cells as in Fig. S1A. Horizontal bars indicate means, error bars mark standard deviation (SD), n=3 biological replicates.
**e.** Flow cytometry of chromatin-bound protein on Wi38 cells synchronized in G0 by contact inhibition in 0.1% FBS for 72 hours and released from G0 into the cell cycle, harvesting 24 hours after release (first cell cycle) and 48 hours after release (second cell cycle). Flow cytometry measured DNA Content (DAPI), Loaded MCM (anti-Mcm2) and DNA Synthesis (EdU). Cells were labeled with 10 μM EdU for 30 minutes before harvesting. Orange cells are S phase $MCM^{DNA}$ positive, blue cells are G1 phase $MCM^{DNA}$ positive, grey cells are $MCM^{DNA}$ negative.
**f.** Loaded MCM in early S phase determined by flow cytometric analysis of early S phase Wi38 from Fig. S1E. Orange line is first cell cycle, grey line is second cell cycle.
**g.** Flow cytometry of chromatin-bound protein on NHF1-htert cells synchronized in G0 by contact inhibition in 0.1% FBS for 72 hours and released from G0 into the cell cycle, harvesting 24 hours after release (first cell cycle) and 48 hours after release (second cell cycle). Flow cytometry measured DNA Content (DAPI), Loaded MCM (anti-Mcm2) and DNA Synthesis (EdU). Orange cells are S phase $MCM^{DNA}$ positive, blue cells are G1 phase $MCM^{DNA}$ positive, grey cells are $MCM^{DNA}$ negative.
**h.** Loaded MCM in early S phase determined by flow cytometric analysis of NHF1 from Fig. S1G. Orange line is first cell cycle, grey line is second cell cycle.
**i.** Flow cytometry of chromatin-bound protein on RPE1 cells synchronized in G0 by starvation in 0% FBS for 72 hours and released from G0 into the cell cycle, harvesting 24 hours after release (first cell cycle) and 48 hours after release (second cell cycle). Flow cytometry measured DNA Content (DAPI), Loaded MCM (anti-Mcm2) and DNA Synthesis (EdU). Orange cells are S phase $MCM^{DNA}$ positive, blue cells are G1 phase $MCM^{DNA}$ positive, grey cells are $MCM^{DNA}$ negative.
**j.** Loaded MCM in early S phase determined by flow cytometric analysis of RPE1 from Fig. S1I. Orange line is first cell cycle, grey line is second cell cycle.

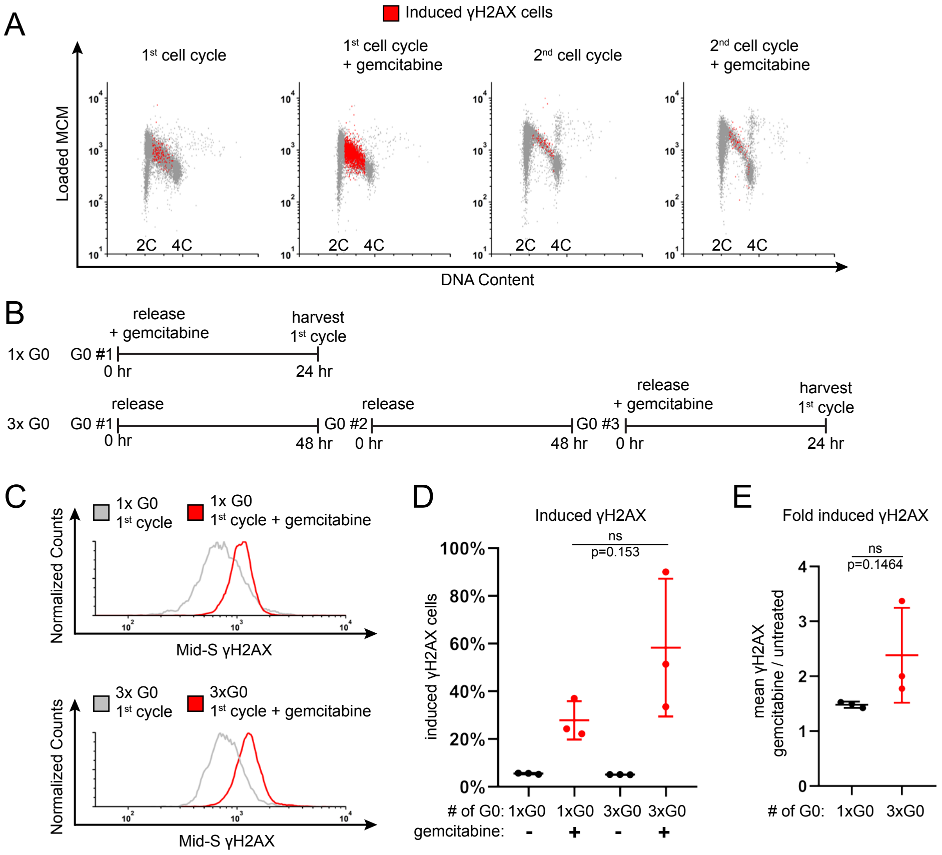


**Supplemental Figure 2. Repeated transitions between G0 and proliferation trend towards an increased replication stress sensitivity**

**a.** Flow cytometry of chromatin-bound protein from cells in Fig. 2B, showing DNA loaded MCM (anti-MCM2). Red cells are gemcitabine (replication stress) induced γH2AX positive, as indicated in Fig. 2C.

**b.** Diagram of experiment. For 1xG0: RPE1 cells were synchronized in G0 by contact inhibition for 48 hours, then released into the cell cycle with 5 nM gemcitabine, harvesting cells 24 hours after release. For 3xG0: RPE1 cells were synchronized in G0 by contact inhibition for 48 hours, released into the cell cycle without gemcitabine and grown to confluency to repeat the G0 by contact inhibition a second time, then repeating once again for a 3rd G0. Then cells were released into the cell cycle with 5 nM gemcitabine, harvesting cells 24 hours after release.

**c.** Flow cytometry of chromatin-bound protein from cells treated as in Fig. S2B, measuring DNA Content (DAPI) and γH2AX (anti H2AX phospho S139). Histograms plot γH2AX of Mid-S phase cells as described in Fig 2C. Upper panel is 1xG0, lower panel is 3xG0. Grey lines are first cell cycle untreated, red lines are first cell cycle treated with 5 nM gemcitabine.

**d.** Percentage of replication stress-induced γH2AX in 1xG0 and 3xG0 release into the first cycle from Fig. S2C. Horizontal bars indicate means, error bars mark standard deviation (SD), n=3 biological replicates. 1xG0 and 3xG0 compared by unpaired, two tailed t test. p=0.153 (ns).

**e.** Comparison of γH2AX intensity per cell presented as fold-change between gemcitabine-treated and untreated cells from Fig. S2C. Horizontal bars indicate means, error bars mark standard deviation (SD), n=3 biological replicates. Samples compared by unpaired, two tailed t test. p=0.1464 (ns).

Figure S3, Matson et al.

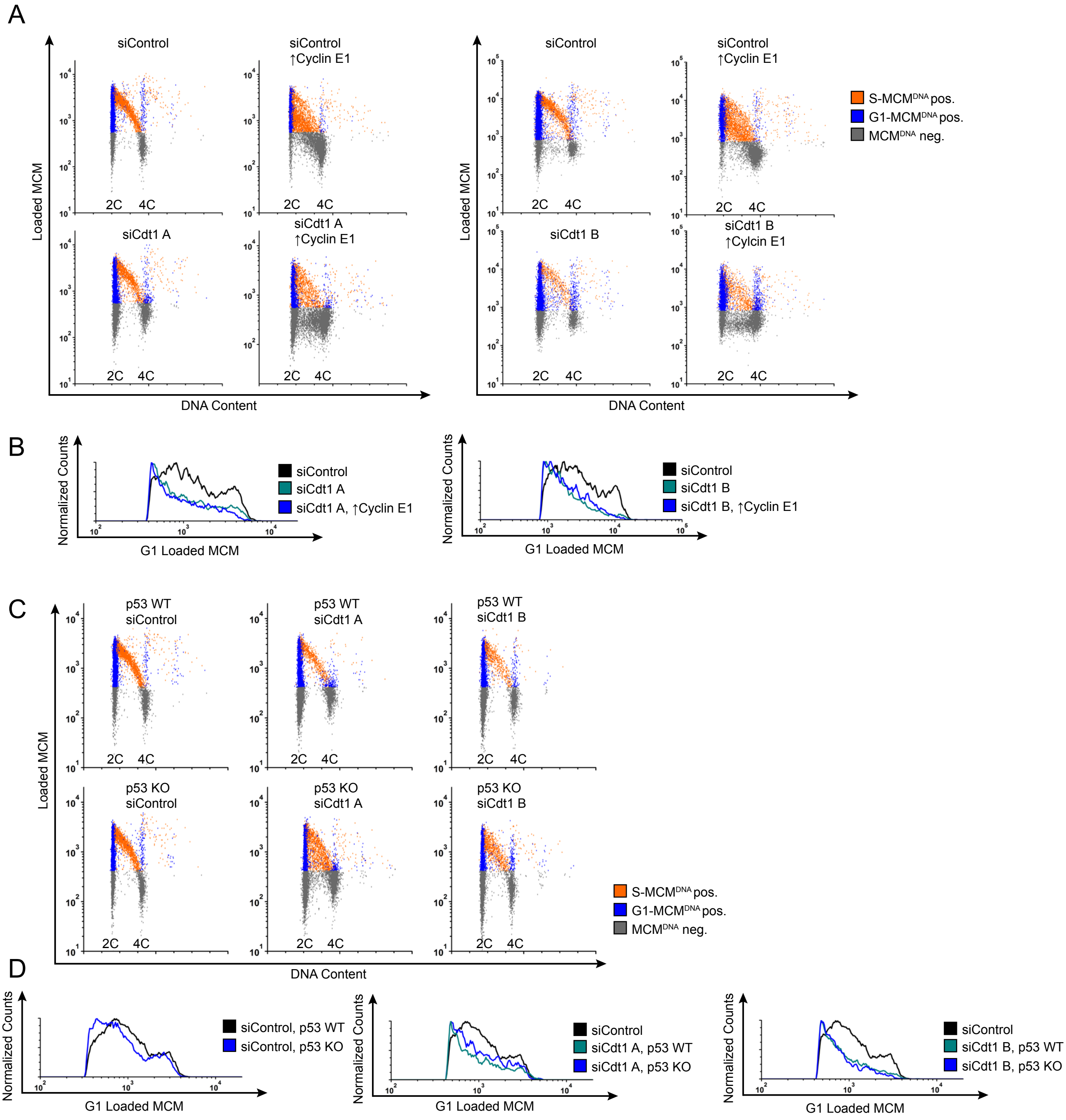

**Supplemental Figure 3. Flow cytometry plots of siRNA in proliferating cells.**
**a.** Flow cytometry of chromatin-bound protein of cells shown in Fig. 3C, measuring DNA Content (DAPI), Loaded MCM (anti-Mcm2) and DNA Synthesis (EdU). Orange cells are S phase $MCM^{DNA}$ positive, blue cells are G1 phase $MCM^{DNA}$ positive, grey cells are $MCM^{DNA}$ negative.
**b.** Loaded MCM in G1 cells from Fig 3C. Black lines are siControl treated cells. Green lines are siCdt1 A (left) or siCdt1 B (right). Blue lines are siCdt1 A, ↑Cyclin E1 (left) or siCdt1 B, ↑Cyclin E1 (right).
**c.** Flow cytometry of chromatin-bound protein of cells shown in Fig. 4C, measuring DNA Content (DAPI), Loaded MCM (anti-Mcm2) and DNA Synthesis (EdU). Orange cells are S phase $MCM^{DNA}$ positive, blue cells are G1 phase $MCM^{DNA}$ positive, grey cells are $MCM^{DNA}$ negative.
**d.** Loaded MCM in G1 cells from Fig. 4C. Left histogram: Black line is siControl, p53 WT, blue line is siControl p53 KO. Middle and right histograms: Black lines are siControl, p53 WT, green lines are siCdt1A, p53 WT (left) or siCdt1 B, p53 WT (right), blue lines are siCdt1 A p53 KO (left) or siCdt1 B p53 KO (right). Note the black siControl p53 WT are the same sample on all three histograms.

Figure S4, Matson et al.

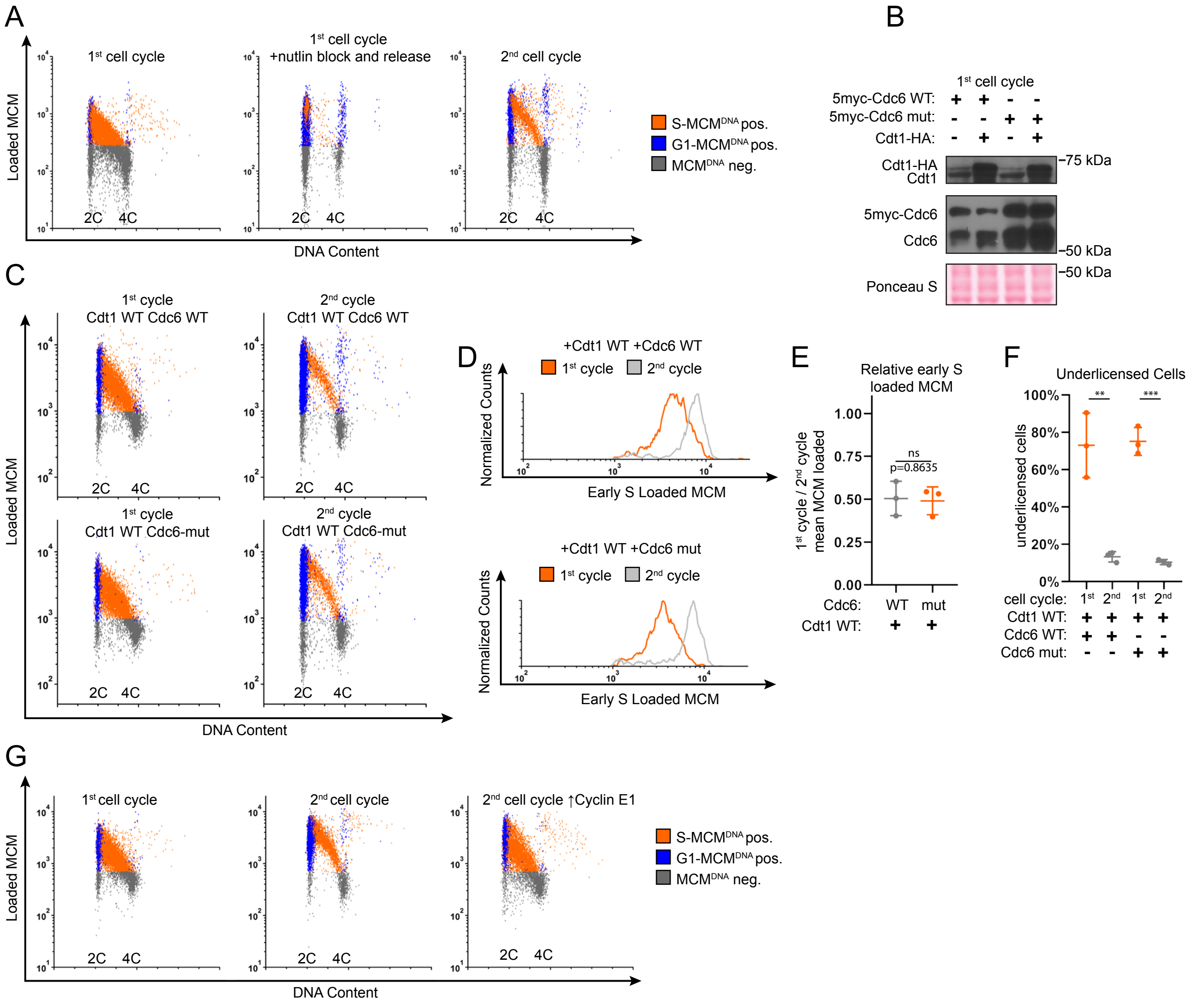

**Supplemental Figure 4. Overproduction of Cdt1 and a stable Cdc6 mutant do not rescue underlicensing in the first cell cycle after G0.**

**a.** Flow cytometry of chromatin-bound protein from cells shown in Fig. 7C, measuring DNA Content (DAPI), Loaded MCM (anti-Mcm2) and DNA Synthesis (EdU). 1st cell cycle is 24 hrs after G0 release, 1st cycle with nutlin block and release are cells treated with 10 μM nutlin-3a from 10-18 hours and released to 26 hours after G0. 2nd cell cycle is 48 hours release from G0. Orange cells are S phase $MCM^{DNA}$ positive, blue cells are G1 phase $MCM^{DNA}$ positive, grey cells are $MCM^{DNA}$ negative.

**b.** Immunoblot for Cdc6 and Cdt1 on total protein lysate from RPE1 cells constitutively producing either 5myc-Cdc6 WT or 5myc-Cdc6-mut (a mutant of Cdc6 that is not targeted for degradation by $APC^{Cdh1}$: R56A, L59A, K81A, E82A, N83A) and a doxycycline inducible Cdt1-HA. RPE1 cells were synchronized in G0 by contact inhibition, treated with 100 ng/mL doxycycline for 4 hours before re-plating cells to release into the first cell cycle, harvesting cells 24 hours after release.

**c.** Flow cytometry of chromatin-bound protein on cells treated as Fig. S4B, harvested 24 hours (first cell cycle) and 48 hours (second cell cycle) after G0 release, measuring DNA Content (DAPI), Loaded MCM (anti-Mcm2), and DNA Synthesis (EdU). Orange cells are S phase $MCM^{DNA}$ positive, blue cells are G1 phase $MCM^{DNA}$ positive, grey cells are $MCM^{DNA}$ negative.

**d.** Loaded MCM of early S phase from Fig. S4C. Orange lines are cells from first cell cycle, grey lines are cells from second cell cyclist. Top panel is RPE1 cells producing Cdt1-HA WT and 5myc-Cdc6 WT, bottom panel is RPE1 cells producing Cdt1-HA WT, 5myc-Cdc6-mut.

**e.** Comparison of early S phase DNA-loaded MCM per cell from Fig. S4D. Values plotted are the ratio of mean loaded MCM of first cell cycle divided by mean loaded MCM of second cell cycle. Horizontal bars indicate means, error bars mark standard deviation (SD), n=3 biological replicates. First and second cell cycles compared by unpaired, two tailed t test. p=0.8635 (ns).

**f.** Percentage of underlicensed cells from early S phase cells in Fig. S4D. Horizontal bars indicate means, error bars mark standard deviation (SD), n=3 biological replicates. First and second cell cycles compared by unpaired, two tailed t test. Cdc6 WT p=0.0041** Cdc6 mut p=0.0001***.

**g.** Flow cytometry of chromatin-bound protein from cells shown in Fig. 7H, measuring DNA Content (DAPI), Loaded MCM (anti-Mcm2) and DNA Synthesis (EdU). Orange cells are S phase $MCM^{DNA}$ positive, blue cells are G1 phase $MCM^{DNA}$ positive, grey cells are $MCM^{DNA}$ negative.
